# Structural basis of norepinephrine recognition and transport inhibition in neurotransmitter transporters

**DOI:** 10.1101/2020.04.30.070219

**Authors:** Shabareesh Pidathala, Aditya Kumar Mallela, Deepthi Joseph, Aravind Penmatsa

**Affiliations:** Molecular Biophysics Unit, Indian Institute of Science, CV Raman Road, Bangalore 560012

**Keywords:** L-Norepinephrine (NE), *Drosophila melanogaster* dopamine transporter (dDAT), dopamine (DA), serotonin (5-HT), solute carrier family, secondary active transport, neurotransmitter sodium symporters, duloxetine, milnacipran, tramadol

## Abstract

Norepinephrine is a biogenic amine neurotransmitter that has widespread effects on cardiovascular tone, alertness and sensation of pain. As a consequence, blockers of norepinephrine uptake have served as vital tools to treat depression and chronic pain. Here, we employ a modified *Drosophila melanogaster* dopamine transporter as a surrogate for the human norepinephrine transporter and determine the X-ray structures of the transporter in its substrate-free and norepinephrine-bound forms. We also report structures of the transporter in complex with inhibitors of chronic pain including duloxetine, milnacipran and a synthetic opioid, tramadol. When compared to dopamine, we observe that norepinephrine binds in a different pose, in the vicinity of subsite C within the primary binding site. Our experiments reveal that this region is the binding site for chronic pain inhibitors and a determinant for norepinephrine-specific reuptake inhibition, thereby providing a paradigm for the design of specific inhibitors for catecholamine neurotransmitter transporters.

**Highlights:**

- X-ray structures of the *Drosophila* dopamine transporter in substrate-free and norepinephrine bound forms.
- Norepinephrine and dopamine bind in distinct conformations within the binding pocket.
- Chronic pain inhibitors S-duloxetine, milnacipran and tramadol bind in the primary binding site and overlap with the norepinephrine-binding pose.
- Selective norepinephrine reuptake inhibition occurs through specific interactions at the subsite C in the primary binding pocket.

## Introduction

Neurotransmitter transporters of the solute carrier 6 (SLC6) family enforce spatiotemporal control of neurotransmitter levels in the synaptic space through Na^+^/Cl^-^-coupled uptake in the central and peripheral nervous systems (Focke et al., 2013; Joseph et al., 2019; Kristensen et al., 2011). Monoamine neurotransmitters affect diverse neurophysiological processes including attention, arousal, sleep, mood, memory, reward, vasodilation and pain (Atzori et al., 2016; Hurlemann et al., 2005; Pertovaara, 2006; Pignatelli and Bonci, 2015; Vatner et al., 1985). Among monoamines, noradrenaline/norepinephrine (NE) is an important neurotransmitter released from the neurons of locus coeruleus in the brain stem that innervate multiple regions of the brain and spinal cord (Sara, 2009). Discovered by vonEuler as a demethylated form of adrenaline (Von Euler, 1946), NE was identified as a neurotransmitter with agonistic effects on the α– and β-adrenergic receptors (Ramos and Arnsten, 2007). The levels of biogenic amines, norepinephrine (NE), dopamine (DA) and serotonin (5-HT), in the neural synapses, are controlled by their cognate transporters, NET, DAT and SERT, respectively (Chang et al., 1996; Kilty et al., 1991; Kristensen et al., 2011; Pacholczyk et al., 1991; Torres et al., 2003). Recent structural studies of the *Drosophila* dopamine transporter (dDAT) and the human serotonin transporter (hSERT) reveal that the SLC6 members closely share their architecture and mechanistic properties (Coleman et al., 2016; Coleman et al., 2019; Penmatsa et al., 2013). The structural similarities among biogenic amine transporters extend their ability to have overlapping substrate specificities particularly between DAT and NET, which are both capable of dopamine and norepinephrine uptake, albeit with varying efficacies (Giros et al., 1994).

Biogenic amine transporters are the primary targets of antidepressants that inhibit monoamine transport and enhance neurotransmitter levels in the synaptic space thereby alleviating depression (Glowinski and Axelrod, 1964; Iversen, 2006). Inhibition of these transporters by psychostimulants like cocaine and amphetamines can result in an acute sensation of reward leading to repeated self-administration and habit-formation (Giros et al., 1996; Iversen, 2000). Specific inhibitors of monoamine uptake including selective serotonin reuptake inhibitors (SSRIs), serotonin-norepinephrine reuptake inhibitors (SNRIs) and norepinephrine reuptake inhibitors (NRIs) have reduced side effects compared to most other antidepressants that non-selectively bind to DAT, NET and SERT (Gether et al., 2006). SNRIs and NRIs are also repositioned and prescribed as medication for chronic pain conditions including neuropathic pain and fibromyalgia (Ashburn and Thor, 2004). They enhance NE levels in the descending pain pathways innervating the dorsal horn of the spinal cord. In this process, NE-mediated activation of the inhibitory α2-adrenergic receptors lead to lowered calcium channel activation and promote hyperpolarization, to reduce chronic pain (Pertovaara, 2006). Most inhibitors of monoamine transport competitively inhibit uptake through interactions in the primary binding site (Apparsundaram et al., 2008; Penmatsa et al., 2013; Wang et al., 2015) although non-competitive inhibition is reported in some instances like citalopram-hSERT interactions (Coleman et al., 2016) and lipid modulators of hGlyTs (Mostyn et al., 2019).

The primary binding site of biogenic amine transporters is divided into subsites A, B and C to delineate the regions of the molecule that interact with substrates and inhibitors (Sorensen et al., 2012; Wang et al., 2013) (Figure 1A). It is also observed that the primary binding site displays remarkable plasticity to accommodate inhibitors of varying sizes (Wang et al., 2015). Alteration of the subsite B residues in dDAT to resemble hDAT or hNET yields a transporter with improved affinities to inhibitors including cocaine, β-CFT and the substrate analogue 3,4-dichlorophenylethylamine (DCP) (Wang et al., 2015). Similarly, the SSRIs also inhibit hSERT through interactions at the primary binding site (Coleman et al., 2016). More recently, the ibogaine-bound structure of hSERT displayed an inward-open conformation, providing crucial insights into the conformational changes associated within the transport cycle of biogenic amine transporters (Coleman et al., 2019).

**Figure 1.**
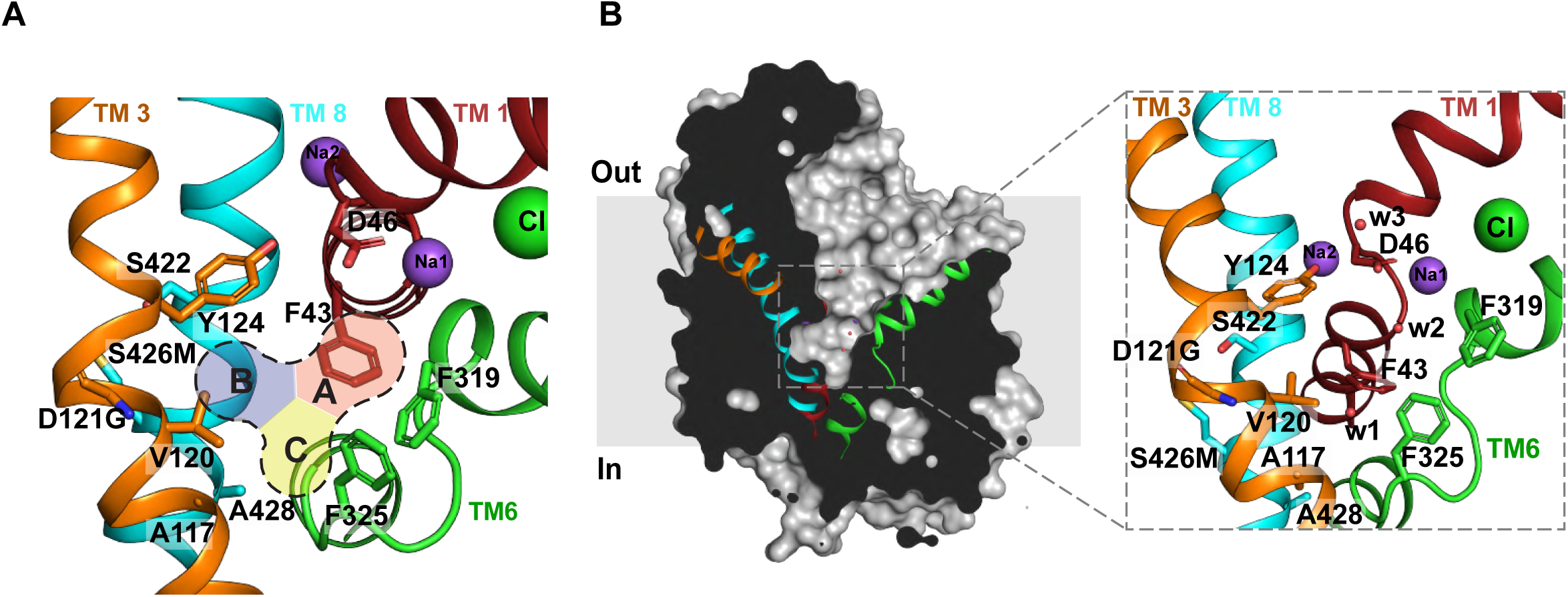
Organization of primary binding site and substrate-free dDAT. **(A)** Close-up view of substrate-free dDAT_subB_ binding pocket, showing the organization of subsites A, B, C of the primary binding site colored as red, blue and yellow, respectively. **(B)** Surface representation of substrate-free dDAT_subB_ structure viewed parallel to the membrane plane. Helices TM1, TM3, TM6 and TM8 are colored as red, orange, green and cyan, respectively. Inset shows the residues lining the primary binding pocket with water molecules indicated as red spheres.

Despite recent progress in understanding the pharmacology and transport mechanism of neurotransmitter transporters through dDAT and hSERT structures, questions linger as to whether dopamine and norepinephrine, both catecholamines, have a similar mode of recognition in hNET. Given the lack of an experimental NET structure, it is also confounding as to how inhibitors can be designed with high specificity towards NET over DAT despite sequence identities greater than 65%. In this context, X-ray structures of dDAT in complex with substrates, including dopamine, 3,4-dichlorophenylethylamine (DCP) and D-amphetamine, have provided a glimpse into substrate recognition and consequent conformational changes that occur in biogenic amine transporters (Wang et al., 2015). Incidentally, dDAT is also capable of NE transport similar to its mammalian orthologues and is well known to have greater affinities towards NE reuptake inhibitors (Penmatsa et al., 2015; Porzgen et al., 2001).

In this study, we employ dDAT as a surrogate of hNET to study the interaction of norepinephrine within the primary binding site. Comparison of different dDAT structures including the substrate-free, dopamine and norepinephrine-bound states allow us to observe and explore interesting differences in substrate recognition in this transporter. Using X-ray structures of dDAT (Table S1) in complex with popularly prescribed inhibitors of chronic pain including S-duloxetine, milnacipran and a synthetic opioid, tramadol, we identify the importance of subsite C as the major determinant of inhibitor specificity between NET and DAT. We also validate these observations through hDAT-like mutagenesis in the subsite C region of dDAT that leads to a loss of affinity towards the NRIs used in the study.

## Results and Discussion

### Modified dDAT mimics hNET primary substrate binding site

The dDAT, much like its human counterpart hDAT, is capable of transporting both dopamine (*K*_M_ 4.5 μM) and norepinephrine (*K*_M_ 55 μM) with varying efficacies and was proposed as a primordial catecholamine transporter in fruit flies (Porzgen et al., 2001) (Figure S1A). The amino acid sequence of dDAT in the primary substrate-binding site has high similarity to hNET and hDAT (Figure S1B), and has pharmacological characteristics closer to hNET whilst having better transport characteristics with dopamine (Porzgen et al., 2001). The dDAT primary binding site is identical to hNET in subsites A and C whereas it differs by two residues in subsite B with polar substitutions; Asp, instead of Gly, at position 121 (149 in hNET) and Ser, instead of a Met, at position 426 (424 in hNET) (Figure S1B, C). Besides these, a vestibule-lining phenylalanine in dDAT was mutated to its hNET counterpart leucine (F471L) to make it resemble hNET. The presence of a leucine at this site was reported to be important for the specific inhibition of hNET by NET-specific χ-conotoxin inhibitor MrIA (Paczkowski et al., 2007). We investigated the effects of substituting these amino acids in the subsite B and vestibule of dDAT on its transport activity. None of the substitutions affected the transport activity of dDAT except for S426M that yielded a transport-inactive form (Figure S1D). Despite the inability of dDAT with these mutations in subsite B to transport catecholamines, it is used in this study as it faithfully reproduces the binding site of hNET, along with the F471L substitution in the vestibule. The construct in the substrate-free, NE-bound and inhibitor-bound forms was crystallized in complex with a heterologously expressed, synthetic version of 9D5 antibody fragment (Fab) that was previously used to crystallize the dDAT (Penmatsa et al., 2013, 2015; Wang et al., 2015) (Figure S2). The NE-bound structures were however determined with both the dDAT_NET_ and the functionally active dDAT_mfc_ constructs carrying the F471L mutation (Figure S1E).

### Substrate-free state is outward-open

The dDAT_subB_ construct with hNET-like mutations was crystallized in the substrate-free Na^+^ and Cl^-^-bound conformation to observe for structural changes in the binding pocket. The transporter structure, determined at 3.3 Å resolution, displays an outward-open conformation with a solvent accessible vestibule that is largely devoid of any specifically bound moieties except for the Na^+^ and Cl^-^ ions at their respective sites (Figure 1B). Despite the absence of bound substrate or inhibitor in the primary binding site, multiple blobs of positive density were observed along the vestibule into which a polyethylene glycol (PEG) was modeled. Within the primary binding site, clear density was observed for most of the residues lining the binding pocket. Solvent accessibility into the primary binding site is unhindered by the F319, which remains splayed open thereby retaining an outward-open conformation (Figure 1B). Interestingly, the side-chain of F325 located in the TM6 linker is positioned in a manner that resembles the antidepressant-bound conformation resulting in a primary binding site with substantial solvent-accessibility (Figure S3). A positive density in the vicinity (∼3.1Å) of F325 was observed into which a water molecule was positioned, allowing the F325 to have lone pair-π interactions (Figure 1B). The outward-open conformation of the dDAT substrate-free state is consistent with the behavior of other NSS members including LeuT whose substrate-free ion-bound conformation is also in the outward-open state (Claxton et al., 2010; Krishnamurthy and Gouaux, 2012). The addition of Na^+^ to LeuT induces opening of the extracellular vestibule suggesting formation of an outward-open state, which is altered upon interactions with substrates like alanine or leucine that induce an occluded state (Zhang et al., 2018). Recent HDX measurements on dDAT and hSERT have clearly indicated the presence of an outward-open conformation in their ion-bound substrate-free state (Moller et al., 2019; Nielsen et al., 2019). Similar observations were evident in all-atom simulations performed on a hDAT model built using dDAT as a template (Cheng and Bahar, 2019). The demonstration of an outward-open conformation in the crystal structure of the substrate-free form of dDAT is a corroboration of these biophysical and computational observations.

### Norepinephrine binds in a different pose in comparison to dopamine

The structural similarity between dopamine and norepinephrine allow them to act as dual substrates for both the NET and DAT orthologues. Whilst the hNET is capable of transporting dopamine (DA) and norepinephrine (NE) with *K*_M_ values of 0.67 μM and 2.6 μM, respectively, the hDAT can transport DA and NE with *K*_M_ values of 2.54 μM and 20 μM, respectively (Giros et al., 1994). Similarly, dDAT is capable of transporting DA and NE with *K*_M_ values of 4.5 μM and 55 μM, respectively (Porzgen et al). It can be consistently observed that both NET and DAT orthologues preferably interact with DA with a higher affinity than NE. It is also observed that in DAT-knockout mice, NET can transport dopamine and substitute for its absence (Moron et al., 2002). The dopamine uptake through dDAT can be competed by L-norepinephrine with a *K*_i_ value of 13.4 μM as observed through whole-cell uptake inhibition studies (Figure 2A).

**Figure 2.**
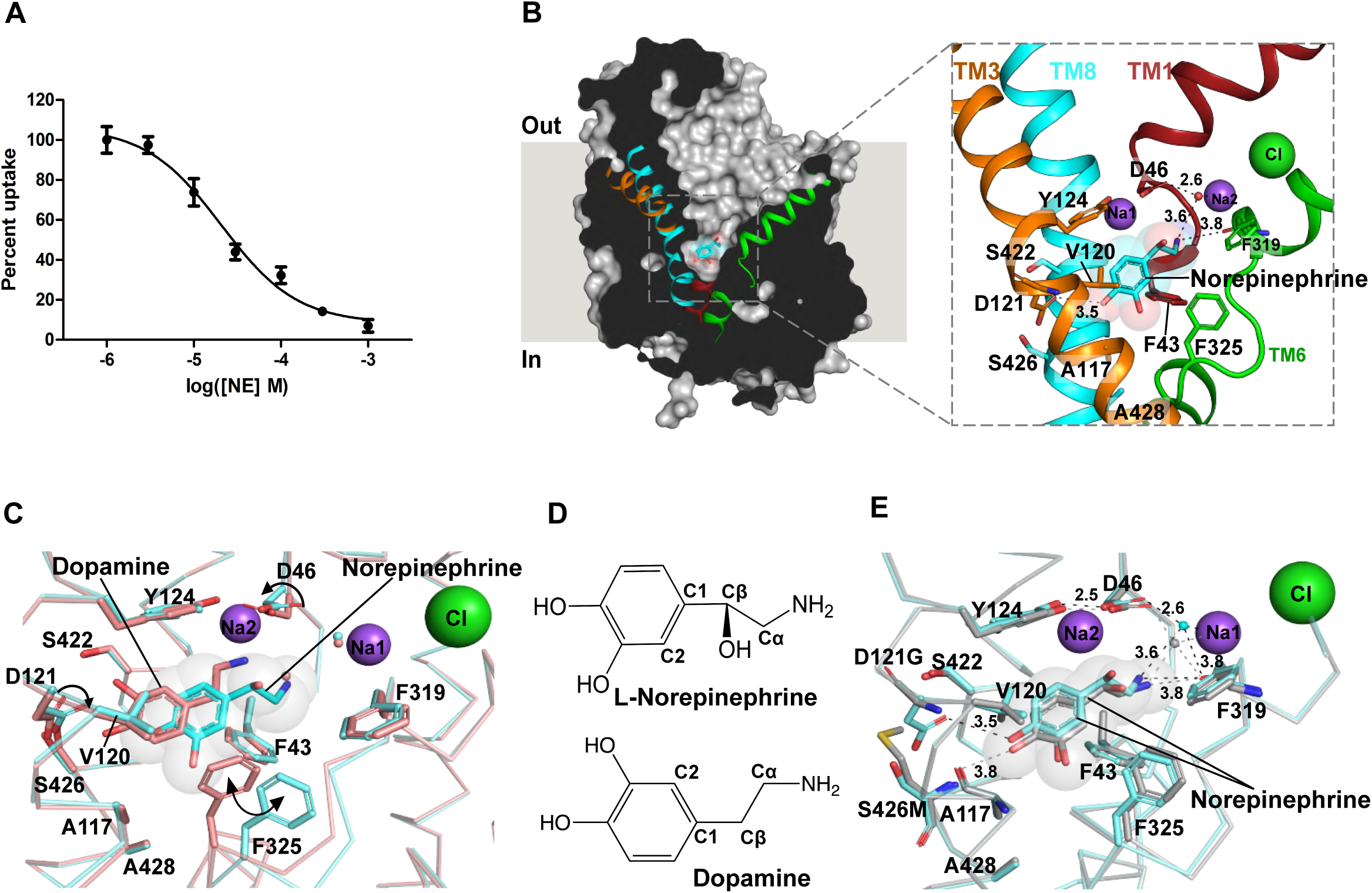
NE binds in a different pose in comparison to DA bound dDAT. **(A)** Inhibition of [^3^H] dopamine uptake by L-norepinephrine with an inhibition constant (*K*_i_) of 13.4 ± 1.4 μM with a functional construct of dDAT. The curve represents one trial of two independent *K*_i_ measurements performed each time in duplicate. **(B)** Longitudinal section of L-norepinephrine (NE)-dDAT_mfc_ complex with L-norepinephrine displayed as cyan spheres. Close-up view of the L-norepinephrine in the primary binding site with surrounding residues displayed as sticks. Hydrogen bond interactions are depicted as dashed lines. **(C)** Comparison of binding pockets of NE-dDAT_mfc_ complex (NE in cyan and backbone in aquamarine) and DA-dDAT_mfc_ (PDB id. 4XP1) complex (DA in deep salmon and backbone in salmon). The D46 side chain remains in position similar to antidepressant-bound structures of dDAT (χ1 torsion angle +85°) unlike dopamine bound state in which D46 side chain rotates to accommodate bound dopamine (χ1 torsion angle −175°). The D121 (TM3) residue in subsite B also shifts (χ2 shifts by 16°) to interact with NE in comparison to the DA bound dDAT structure. The position of F325 shifts by nearly 2Å (Cβ) with a corresponding rotation of the phenyl group by 51° (χ2 torsion angle CD1-Cγ–Cβ-Cα) (**D)** Chemical structures of L-norepinephrine and dopamine represented in their bound conformations. **(E)** Superposition of binding pockets of the NE-dDAT_mfc_ structure (NE and backbone in cyan) with the binding pocket of the NE-dDAT_NET_ structure carrying hNET-like mutations in subsite B (NE and backbone in gray). NE was modeled into densities at near identical positions in both the structures.

The norepinephrine-bound dDAT_mfc_ structure reveals clear density for NE bound within the primary binding site of the transporter (Figure 2B, S4A). The primary amine of NE interacts in the subsite A region forming hydrogen bonds with carbonyl oxygens of F43 and F319 main chain and the carboxylate side chain of D46 via a water molecule (Figure 2B). The D46 residue in the DA-bound dDAT structure undergoes a χ1 torsion angle shift of 100° relative to that of the NE-bound structure to interact with the primary amine of DA. However, no such shift was observed in the NE-bound DAT structure (Figure 2C). The primary amine interacts with a Na^+^-coordinating water molecule akin to the D-amphetamine and DCP-bound structures (Figure S5). Interestingly, the binding pose of NE does not resemble that of DA in the binding pocket despite both the substrates being catecholamines (Figure 2C, D). Earlier computational studies predicted that the catechol group of NE predominantly interacts with subsite B region (Koldso et al., 2013; Schlessinger et al., 2011). However, we observe that the NE catechol group binds in the vicinity of subsite C in the region between TM6 linker and TM3 with no conformational changes in the binding pocket in comparison to the outward-open substrate-free form of dDAT_subB_ (Figure 2C, S5). The binding of NE in the primary binding site resembles a lock-and-key association in comparison to the induced-fit interaction observed with dopamine binding. Clear density for the β-OH group of NE is observed in the primary binding site (Figure S4) and the β-OH group faces the solvent accessible vestibule. The para-OH group of the catechol ring retains interactions with the side chain carboxyl of D121 that undergoes a rotation of 16° along the χ2 torsion angle in comparison with the DA-bound structure, to facilitate interactions with NE (Figure 2C). The meta-OH group of NE displaces the water molecule observed in the substrate-free state, fitting snugly into the gap between A117 and TM6 linker adjacent to the F325 side chain (Figure S5A). Dopamine, on the other hand, interacts closely with residues in subsite B, displaying a near 180° flip in the position of the catechol group, relative to NE. This induces a shift in the position of F325 to facilitate edge-to-face aromatic interactions with DA, which are absent in the NE-bound structure (Figure 2C).

The subsite B residues in dDAT differ from hNET and hDAT at two positions, D121G (TM3) and S426M (TM8) (Figure S1B, S1C). In order to evaluate whether hNET-like substitutions in the subsite B of dDAT would influence and shift the conformation of NE to DA-like pose, we crystallized NE in complex with dDAT_NET_ construct that has D121G and S426M substitutions. Despite hNET-like substitutions, no major shifts in the position of NE were observed in the binding site. Despite lack of the D121 sidechain, the para-OH group of NE establishes interactions with the main chain carbonyl oxygen of A117 (Figure 2E).

The comparison between DA and NE-bound crystal structures show substantial difference between their preferred conformers within the binding pocket. The catechol ring of NE is positioned in an opposite orientation relative to that of DA with a slight difference in the C_2_C_1_-C_β_C_α_ torsion angle (26°) (Figure 2D). This difference in their binding poses, despite being very similar catecholamines, can be attributed to the presence of β-OH group that is known to restrict the conformational freedom of NE along the C_1_-C_β_ bond. Between the stable conformers of NE, the energy barrier for the catechol rotation along C_1_-C_β_ bond is nearly 9-12 kcal/mol whereas in the case of DA it is as low as 0.3-0.6 kcal/mol as shown in simulations performed on both the neurotransmitters (Nagy et al., 1999; Nagy et al., 2003). The large energy barrier between NE and DA likely restrains NE to a fixed conformation leading it to binding closer to the subsite C of the binding pocket; whereas DA, owing to the greater flexibility, binds in proximity to subsite B. The induced-fit conformational change in response to dopamine binding and the lack thereof in NE is the likely reason for both DAT and NET to have a greater propensity to interact with DA in comparison to NE.

### S-duloxetine and Milnacipran are competitive inhibitors of norepinephrine transport

The ability of serotonin-norepinephrine reuptake inhibitors (SNRIs) to alleviate chronic pain by blocking NET activity in the descending pain pathways has allowed drugs like S-duloxetine and milnacipran to be repositioned for treatment of neuropathic pain and fibromyalgia (Arnold et al., 2007). Much like the other inhibitors/antidepressants characterized including nortriptyline, nisoxetine and reboxetine in complex with dDAT (Penmatsa et al., 2013, 2015) and paroxetine, S-citalopram, fluoxetine in complex with hSERT (Coleman and Gouaux, 2018), both S-duloxetine and milnacipran interact at the primary binding site of the dDAT structure (Figures 3, 4). The electron densities for both drugs are unambiguous and conform to the general principles of inhibitor interactions with the transporter (Figure S4C, D).

**Figure 3.**
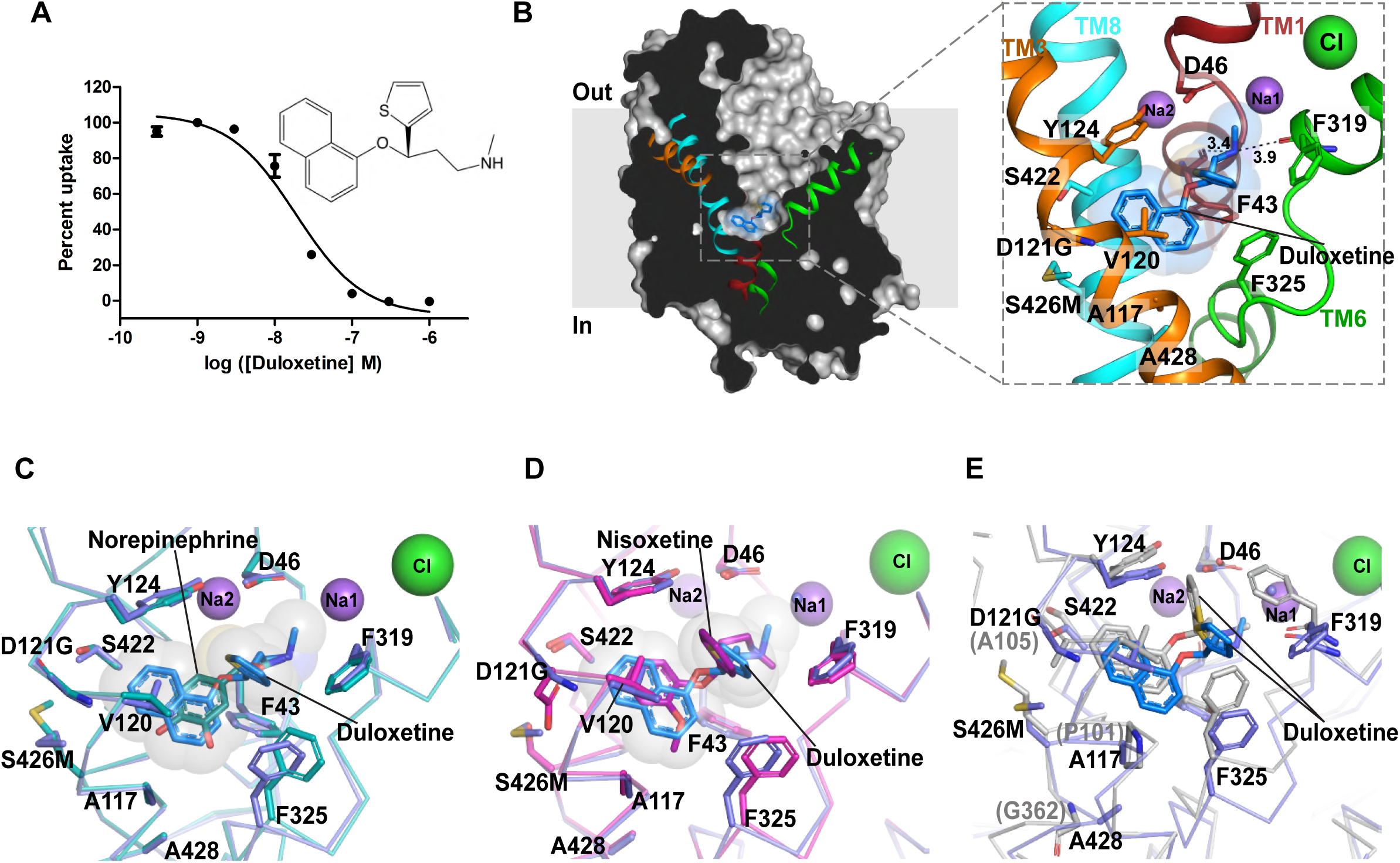
S-duloxetine binds in the primary binding site. **(A)** Inhibition of [^3^H] dopamine uptake with S-duloxetine (*K*_i_ = 14.3 ± 7.4 nM). The graph is representative of one of three trials each performed in duplicate. Error bars represent the range of the [^3^H] dopamine uptake calculated as percent inhibition. **(B)** Surface representation of dDAT_subB_ with bound S-duloxetine shown as blue sticks and transparent spheres in the primary binding site. Inset shows orientation of the inhibitor in the binding pocket with residues in the vicinity represented as sticks. **(C)** Binding site comparison of S-duloxetine with NE bound dDAT_mfc_. The phenyl group of F325 shifts by 40° to retain edge-to-face aromatic interactions with the naphthalene group of S-duloxetine. **(D)** Overlay of the dDAT bound to S-duloxetine and nisoxetine (magenta) (PDB id. 4XNU). The surrounding residues display an identical binding pose between the two structures. **(E)** Overlay of duloxetine bound dDAT_subB_ and LeuBAT structures (PDB. id. 4MMD) displaying similar pose of the inhibitor in both the structures. LeuBAT displays a prominent occluded conformation whereas dDAT retains an outward-open conformation.

**Figure 4.**
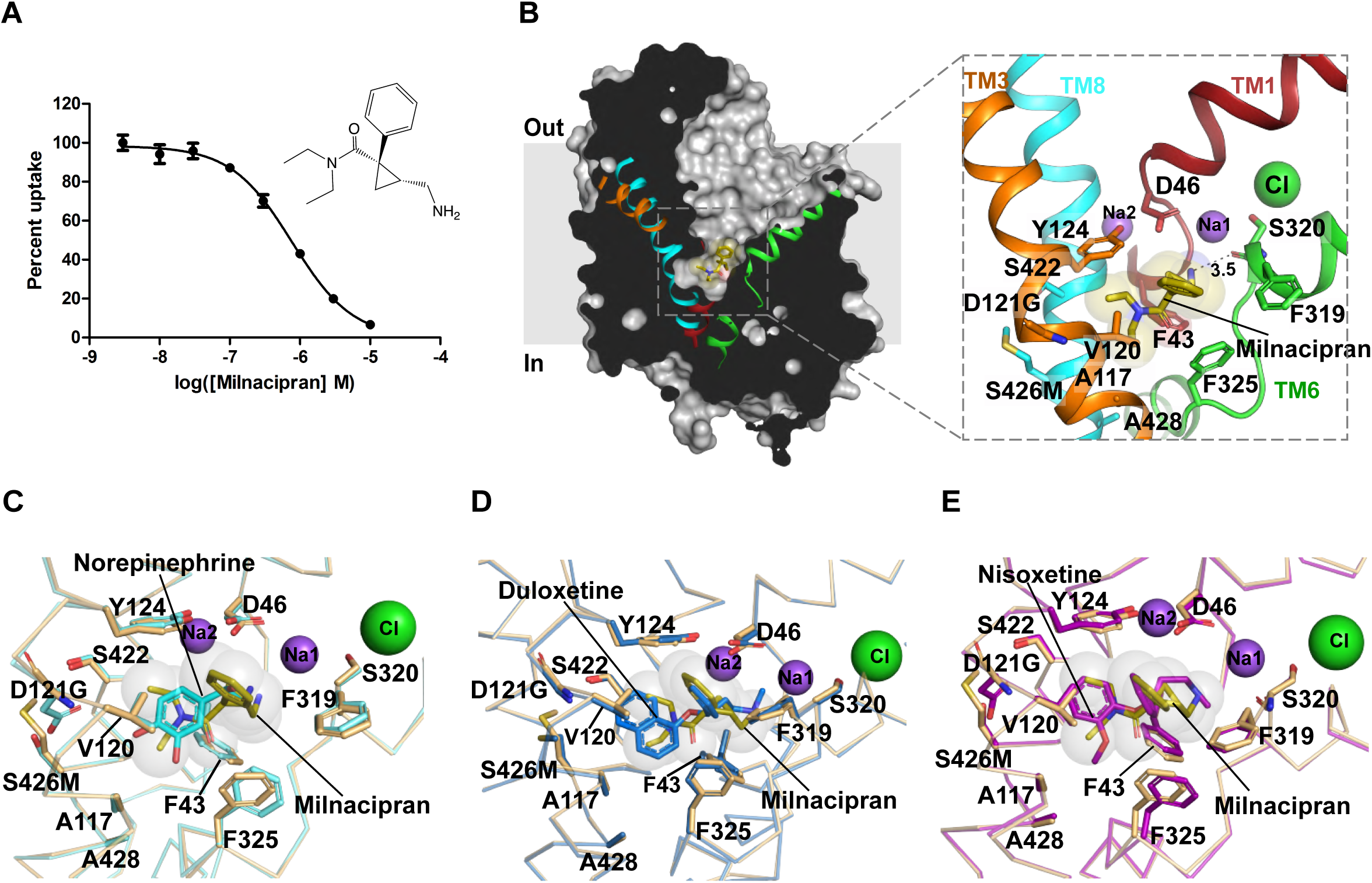
Milnacipran is a competitive inhibitor of NE uptake. **(A)** [^3^H] dopamine transport inhibition by increasing concentrations of 1R-2S milnacipran with a *K*_*i*_ value of 584 ± 71 nM. Graph represents one of two repetitions of the experiment, each performed in duplicate. Error bars represent the measured range of percent inhibition at individual concentrations. Inset displays the chemical structure of 1R-2S milnacipran. **(B)** Longitudinal section of dDAT_NET_ in complex with milnacipran bound (olive sticks) in the primary binding site with residues in proximity displayed as sticks. **(C)** Milnacipran binding pose compared to NE binding in dDAT_mfc_ reveals an angular shift of F325 by 35° to retain hydrophobic interactions with the diethyl group of milnacipran. **(D)** Milnacipran bound dDAT_NET_ vs duloxetine bound dDAT_subB_ display identical positions of the surrounding residues. **(E)** Structural overlap of nisoxetine bound dDAT (PDB id. 4XNU) with milnacipran bound dDAT_subB_ revealing the surrounding residues in identical binding positions.

S-duloxetine, owing to its large surface area (surface area 753.7 Å^2^) exhibits maximal occupancy of the primary binding pocket. The drug inhibits dopamine uptake with a *K*_i_ value of ∼14 nM (Figure 3A), consistent with the *K*_i_ values observed for hNET inhibition by duloxetine (Bymaster et al., 2003). The high affinity is an outcome of its ability to snugly fit into the cavernous primary binding site of NSS transporters. In duloxetine, the propanamine group interacts with the main chain carbonyl oxygens from residues F43 and F319 with the D46 residue in the vicinity (Figure 3B). The secondary amine can also mediate π – cation interaction with the side chain of F43. The naphthyloxy ring interactions in the binding pocket extend from subsite B to subsite C, wedging into space sculpted by residues including Y124, D121G, S426M, V120, A117 in TM3 and TM8 followed by edge-to-face aromatic interactions with F325 in subsite C. The thiophene ring is positioned with some elevation within the binding pocket to sterically block the closure of the F319 and thus precluding the formation of occluded state during transport (Figure 3B). The duloxetine position in the binding pocket overlaps with NE and nisoxetine binding to a large extent with the naphthalene ring taking the place of the methoxyphenoxy ring of nisoxetine, a specific inhibitor of NE reuptake (Figure 3C, D). Similarly, when compared with the cocaine bound structure of dDAT, one of the aromatic groups of the naphthalene ring in duloxetine overlaps with the benzoyl moiety of cocaine (Figure S6). However, lack of an additional hydrophobic moiety in cocaine makes it a moderate inhibitor of NE uptake (Hoepping et al., 2000), relative to duloxetine. The position of duloxetine in the binding pocket is very similar to the LeuBAT-duloxetine complex elucidated earlier (Wang et al., 2013), thus corroborating LeuBAT as a relevant model system to study pharmacology of biogenic amine transporters (Figure 3E).

Milnacipran has an unconventional structure with a cyclopropyl skeleton having both a primary amine and tertiary amine (N, N-diethyl) being part of the drug structure (Andersen et al., 2009). The drug lacks large aromatic moieties that are commonly observed with most NSS inhibitors. Milnacipran inhibits dopamine transport by dDAT with a *K*_i_ value of 0.58 µM, which is much lower than S-duloxetine (Fig 4A). Like duloxetine, milnacipran also binds in the primary binding site and overlays well with NE (Figure 4B, C). The primary amine of aminomethyl group in milnacipran interacts with the side-chain of D46 and the main-chain carbonyls of F43 and S320 in the subsite A by hydrogen bonds (Figure 4B). The phenyl group attached to the chiral centre at the cyclopropane group overlaps with the thiophene group of duloxetine and phenyl ring of nisoxetine (Figure 4D, E). Interestingly, the N, N diethyl group, which is usually occupied by bulky aromatic groups in most of the inhibitors, does not wedge deeply into the subsite B as observed with cocaine (Figure S6C) and retains hydrophobic interactions in the vicinity of subsite C. The absence of a bulky aromatic group wedging into subsite B could be the reason for the lowered transport inhibition observed in milnacipran in comparison to duloxetine. Indicative of this is an observation that altering the hydrophobicity of the binding site by substituting residue V148 (V120 in dDAT_NET_) in hNET to an isoleucine led to a 17-fold enhancement of milnacipran’s inhibitory potency (*K*_i_) (Sorensen et al., 2012).

### Synthetic opioid tramadol blocks transport by interacting with subsite C

It is well known that some synthetic opioids have a dual mechanism of action for pain relief by serving as agonists of μ-opioid receptors and blockers of NE and 5-HT uptake. Tramadol is a popularly used synthetic opioid with a dual ability of activating opioid and NE based analgesic pathways (Grond and Sablotzki, 2004). While tramadol is considered a safe drug, it induces opioid-like symptoms and dependence when used in supra-therapeutic doses (Dayer et al., 1997). The demethylated metabolite of tramadol, O-desmethyl tramadol (desmetramadol) is a better agonist for opioid receptor whilst tramadol, particularly S,S-tramadol is specific to norepinephrine transport inhibition (Rickli et al., 2018). Tramadol and desmetramadol have structural resemblance to the antidepressant, desvenlafaxine (Figure 5A). Tramadol inhibits the transport activity of dDAT with a *K*_i_ of 7.1 µM (Figure 5), which is substantially lower than that of S-duloxetine and milnacipran. Similarly, tramadol has lower *K*_*i*_ value of 2.6 μM relative to the other studied inhibitors for competitively displacing nisoxetine from the binding pocket (Figure S7). Despite weaker affinities to the transporter, tramadol is highly specific to NET/SERT over DAT displaying nearly fifty-fold greater affinity to NET over DAT (Rickli et al., 2018). The structure of the tramadol-DAT_NET_ complex reveals that the drug binds to the primary binding site with the tertiary amine of the dimethyl amino group interacting with subsite A residues D46 and carbonyl oxygens of F43 and F319 (Figure 5B). The 1-cyclohexanol group takes a similar position to the thiophene group of S-duloxetine and phenyl groups of milnacipran and nisoxetine to sterically prevent the formation of an outward-occluded conformation in the transport process (Figure S3). The lower affinity of tramadol to inhibit neurotransmitter uptake compared to other SNRIs could be attributed to the lack of an aromatic moiety in the close vicinity of the subsite B. This distinction is even more apparent when the tramadol bound dDAT_NET_ structure is compared to cocaine bound dDAT_mfc_ structure where the benzoyl group of cocaine is clearly wedged into subsite B in comparison to tramadol’s methoxy phenyl ring that is primarily in the vicinity of subsite C (Figure 5E). The methoxy phenyl ring interacts with the side chain of Y124 by aromatic edge-to-face interactions and fits into the hydrophobic pocket lined by the side-chains of A117, V120, A428 and F325 of subsite C. The position of methoxyphenyl group coincides very closely with the catechol group of NE and methoxy group overlaps well with the meta-OH of NE (Figure 5D). Much like nisoxetine and reboxetine, the methoxy group of tramadol occupies the space between A117 and F325 and interestingly, the demethylated metabolite of tramadol, desmetramadol is a weaker inhibitor of hNET (IC_50_ = 6 μM) relative to tramadol (IC_50_ = 2 μM) indicating the importance of this hydrophobic interaction in enhancing the efficacy of selective NE reuptake inhibition (Rickli et al., 2018). The lack of interactions in subsite B and the presence of substitutions in subsite C (A117S and A428S) unique to human DAT, makes tramadol a weak hDAT inhibitor with a IC_50_ of ∼100 μM (Rickli et al., 2018). This is evident from the similar *K*_*i*_ values of tramadol in displacing nisoxetine bound to dDAT_mfc_ and dDAT_NET_ proteins wherein dDAT_NET_ has hNET-like D121G/S426M substitutions in subsite B (Figure S7). In contrast, inhibitors like cocaine, which primarily bind to subsite B have a 10-fold increment in their ability to compete for nisoxetine in the presence of hNET-like mutations in subsite B (Wang et al., 2015). The observations from tramadol-dDAT complex clearly indicate that NET-specific inhibition occurs through interactions at subsite C. It is also evident that the lack of interactions at subsite B compromises the affinity of tramadol, but clearly fails to influence the specificity of NET inhibition in comparison to DAT (Rickli et al., 2018).

**Figure 5.**
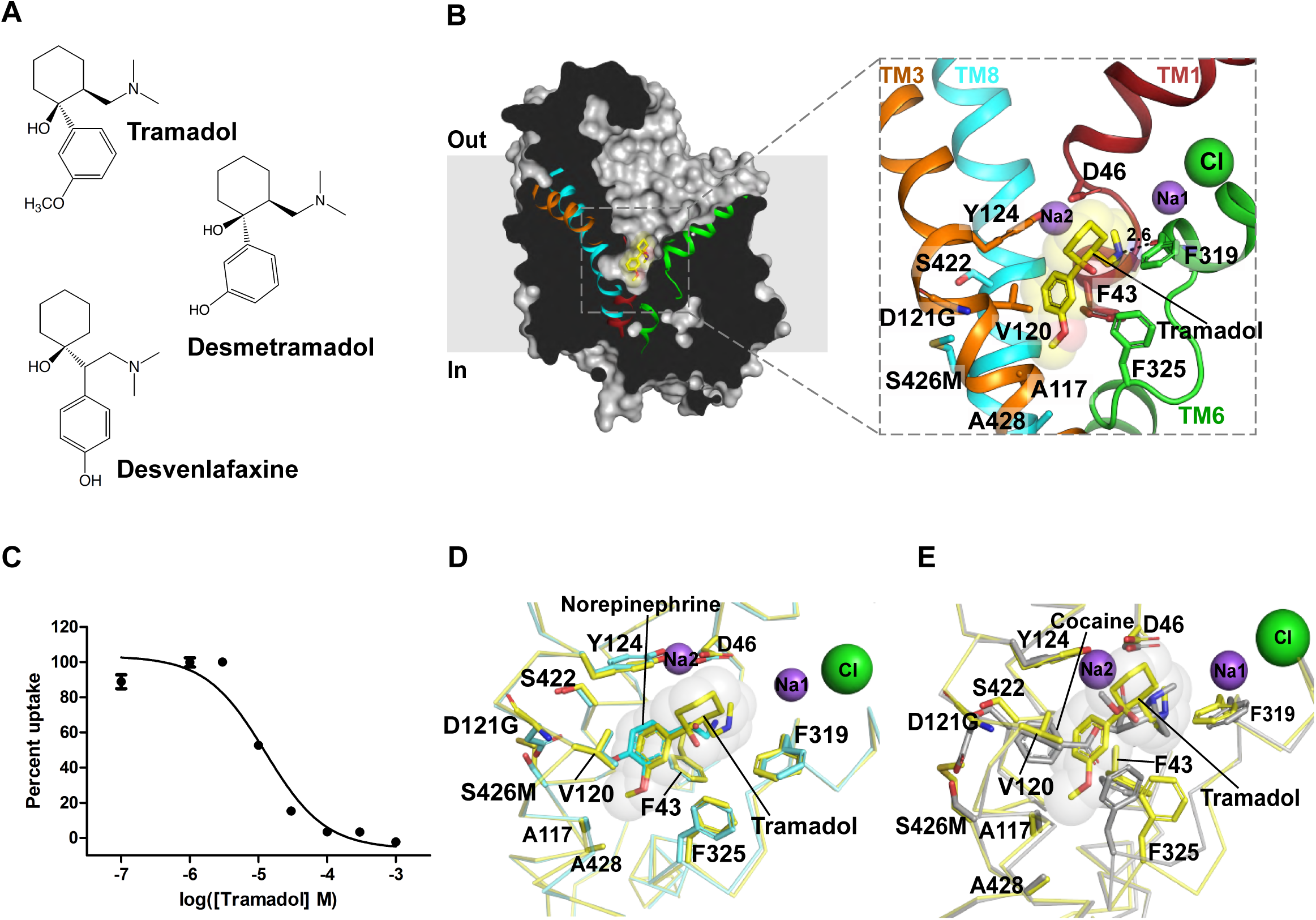
Tramadol is a synthetic opioid that inhibits NE uptake. **(A)** Chemical structures of inhibitors, tramadol, desmetramadol and desvenlafaxine. **(B)** Structure of tramadol (yellow) bound dDAT_NET_ viewed parallel to the membrane plane where tramadol was observed in the primary binding site. **(C)** [^3^H] dopamine uptake inhibition observed with increasing concentrations of tramadol displaying a *K*_i_ value of 7.1 ± 1.6 μM. The graph is a representative of three repetitions, each performed in duplicate. **(D)** Structural comparison around the primary binding site between tramadol bound dDAT_NET_ and NE bound dDAT_mfc_ wherein the methoxyphenyl group of tramadol overlaps exactly with the catechol ring of NE. No differences in the positions of the binding site residues were observed. **(E)** Comparison of tramadol bound dDAT_NET_ structure with cocaine bound dDAT_mfc_ structure (PDB id. 4XP4). The benzoyl ester group of cocaine clearly interacts with subsite B whereas tramadol’s aromatic group interacts primarily at subsite C, within the primary binding site.

### Ligand binding to subsite C influences specificity of NE uptake inhibition

The structures of SNRIs duloxetine, milnacipran and tramadol in complex with the dDAT show a progressively smaller aromatic moiety that interacts with the subsite B and C in the primary binding pocket of dDAT. In an earlier study, it was observed that the aromatic moieties in drugs like cocaine, RTI-55 and nisoxetine interact closely in the subsite B and have enhanced affinities when hNET-like substitutions D121G and S426M are made in the pocket (Wang et al., 2015). Interestingly, this improvement in affinity is not apparent in the case of SNRIs employed in the study where the *K*_i_ values remain unchanged or weaken when substitutions in subsite B are made to improve the identity of dDAT binding pocket to hNET (Figure S7). The minimal effect of subsite B substitutions on the affinity of duloxetine, milnacipran and tramadol suggest that determinants of NET specificity lie elsewhere in the binding pocket. Earlier studies have posited that non-conserved residues in the primary binding site are responsible for selective inhibition of biogenic amine uptake (Andersen et al., 2015; Andersen et al., 2011). A close examination of the binding pocket reveals that hDAT and hNET have subtle differences within them. The dDAT, while largely resembling hNET in the binding pocket differs from hDAT in the residues A117(TM3), A428(TM8) and S422(TM8). The former two residues line subsite C in the vicinity of the TM6 linker where NET specific drugs like nisoxetine, reboxetine, milnacipran and tramadol interact. In order to evaluate the role of these three residues on the ability of drugs to inhibit neurotransmitter transport, hDAT-like mutations A117S, A428S, S422A and a combination of A117S/A428S were introduced into the functional construct of dDAT. Effects of these mutations were analyzed through uptake inhibition using the three SNRIs employed in this study (Figure 6A-C). All the three drugs duloxetine, milnacipran and tramadol display no effect in IC_50_ values with the S422A mutant and retained near WT-like inhibition (Figure 6D). Individual substitution of A428S at subsite C caused a marginal (2-5 fold) loss of uptake inhibition, whereas the A117S mutation did not cause any significant change (Figure 6D). However, a combination of A428S and A117S caused a substantial loss in the ability of duloxetine, milnacipran and tramadol to inhibit dopamine transport as observed by a 5, 7 and 15-fold increase in their IC_50_ values, respectively (Figure 6D). The IC_50_ values for tramadol and milnacipran obtained with the A117S-A428S double mutant are very similar to the reported IC_50_ values of these inhibitors in hDAT (Chen et al., 2008; Rickli et al., 2018). Thus, these substitutions clearly indicate the importance of subsite C region in dictating the specificity of individual inhibitors that exploit this hydrophobic cavity to gain NET specificity. The polar substitutions observed in this region in hDAT lead to a reduced ability of SNRIs with hydrophobic moieties to bind efficiently, thereby compromising their ability to interact with the hDAT (Figure 7A, B).

**Figure 6.**
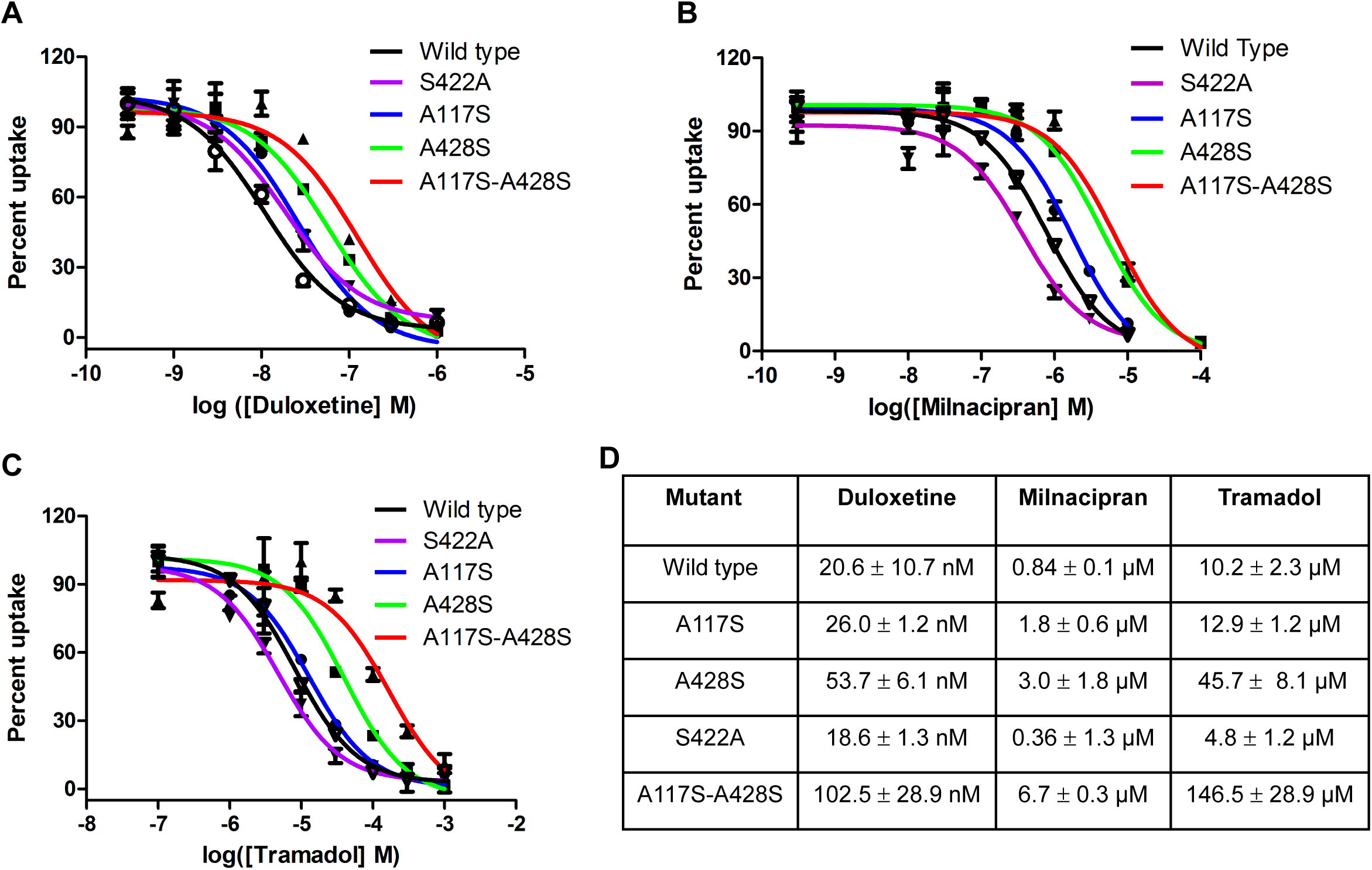
hDAT-like mutagenesis of residues leads to reduced affinities of NET specific inhibitors. **(A)** Dose-response curves of [^3^H] dopamine uptake inhibition by duloxetine with a functional construct of dDAT carrying hDAT-like mutations in the primary binding site including wild-type (black), S422A (purple), A117S (blue), A428S (green) and A117S/A428S double mutant (red). **(B), (C)** Similar uptake inhibition curves plotted using milnacipran and tramadol, respectively. All the graphs represent one of two independent repetitions performed in duplicate. **(D)** Table indicates IC_50_ values of individual mutant inhibition curves with all three inhibitors. Substitution of subsite C residues A428S and double mutant A117S and A428S display maximal loss of affinity for SNRIs duloxetine, milnacipran and tramadol.

**Figure 7.**
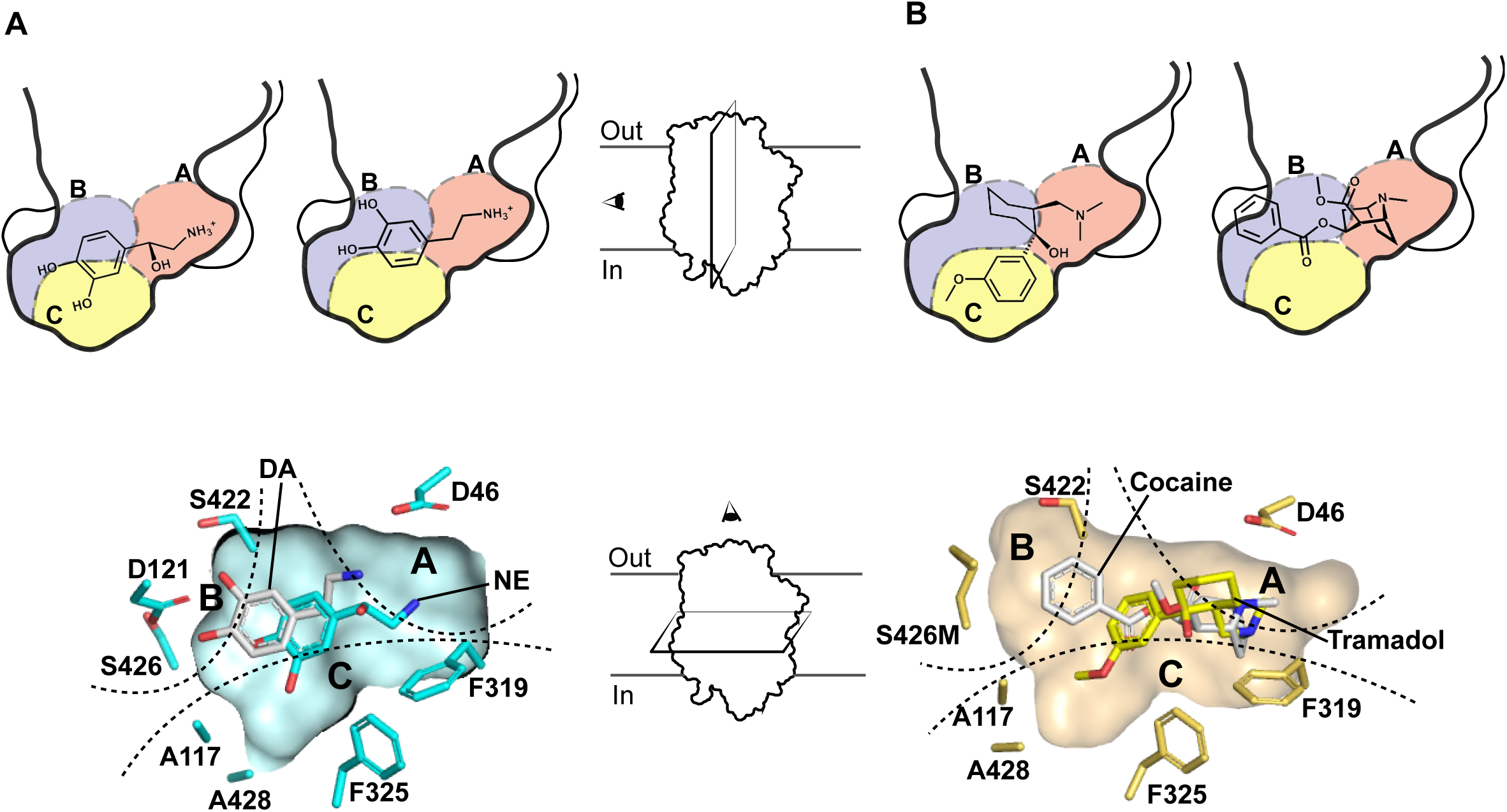
Distinct binding poses of substrates and NET inhibitors. **(A)** Lateral sections of the primary binding pocket comparisons between NE and DA bound states reveal distinct conformations and binding poses of the two substrates in the subsite C and subsite B, respectively (top). Transverse section of the binding pocket clearly reveal similar distinctions between NE and DA (bottom). **(B)** Lateral section comparisons between tramadol and cocaine, displaying primary interactions of their aromatic moieties in subsite C and subsite B respectively (top). Transverse section of the binding pocket showing tramadol overlapped with cocaine indicates clear differences in interaction of a non-specific inhibitor, cocaine towards subsite B and an SNRI, tramadol interacting preferentially with subsite C.

## Conclusions

This study, using X-ray structures of dDAT in complex with NE and NET – specific inhibitors, highlights the discrepancies in catecholamine recognition in neurotransmitter transporters and explores the basis for NET-specific reuptake inhibition over dopamine reuptake inhibition, despite the close similarity between NET and DAT. The catechol group of NE is observed to interact primarily at subsite C in the vicinity of NET-specific residues A117 and A428. The binding of NE displaces water molecules in the binding pocket observed in the substrate-free state and does not induce any local conformational changes in the binding, contrary to DA (Figure 7A). DA was previously observed to interact closely with its catechol group in the vicinity of subsite B leading to shifts in the positions of residues D46 and F325 to retain interactions with the neurotransmitter (Wang et al., 2015). The absence of these changes in NE-dDAT complex could be attributed to the reduced flexibility of NE, in comparison to DA, due to the presence of the β-OH group in its structure (Nagy et al., 2003).

The NET-specific chronic pain inhibitors, duloxetine, milnacipran and tramadol, compete for NE binding site through aromatic groups. These aromatic moieties can interact and snugly fit at subsites B and C to retain high affinity and selectivity towards specific biogenic amine transporters. However, hDAT-like mutations within subsite C of dDAT, A117S and A428S alter the polarity of the binding pocket and weaken the hydrophobic interactions and prevent functional groups like the methoxyphenyl group of tramadol from accessing the subsite through steric block (Figure, 7B). On the other hand, non-specific inhibitors of biogenic amine transport, for instance cocaine, primarily interact with subsite B wherein hNET-like mutations D121G and S426M in dDAT enhance cocaine affinity by 10-fold. The primary interactions of cocaine at subsite B induces a plastic reorganization of the binding pocket as the subsite C residue F325 compensates for lack of bulky aromatic group in cocaine through local conformational changes to establish aromatic π-stacking interactions (Wang et al., 2015). Through our results, we infer that NET-specific inhibitors could be designed with primary interactions at subsite C whilst non-specific high affinity interactions are observed in inhibitors that interact at subsite B. Taken together, the results of this work convey the unique facets of catecholamine recognition within the same binding pocket and establish the roles of individual subsites in dictating inhibitor selectivity and affinity among biogenic amine transporters. The findings can effectively be used for selective inhibitor design targeting pharmacological niches as widespread as depression and chronic pain.

## Author contributions

SP performed protein expression, purification, crystallization and uptake measurements. AKM developed the methodology for heterologous expression of the antibody fragment and performed biochemical assays involving binding. DJ performed biochemical analyses and aided in the preparation of figures. AP designed the study and performed crystallographic analyses. AP wrote the manuscript with inputs from all the authors.

## Acknowledgements

The authors would like to thank all members of the Penmatsa lab for suggestions. The authors would like to thank Dr. Eric Gouaux, Vollum Institute, OHSU for the gift of the dDAT pEG-BacMam constructs. We thank Dr. Vinothkumar KR at the National Centre for Biological Sciences, Bangalore for comments on the manuscript. The authors would like to thank the staff of northeastern collaborative access team (NECAT), Advanced Photon Source, particularly Dr. Surajit Banerjee for access and help with data collection. We thank the beamline staff at the Elettra XRD2 particularly Dr. Babu Manjashetty and Dr. Annie Heroux for beamline support. Access to the XRD2 beamline at Elettra synchrotron, Trieste was made possible through grant-in-aid from the Department of Science and Technology, India, vide grant number DSTO-1668. We acknowledge ESRF access program of the RCB (Grant # BT/INF/22/SP22660/2017) of the Department of Biotechnology, India. We would like to thank the staff of PX beamline of the Swiss Light Source for beamline access and support.

## Experimental Procedures

### List of constructs

The *Drosophila melanogaster* dopamine transporter construct used for performing transport assays has a deletion of 20 amino acids in the amino terminal (Δ1-20) and a deletion in the extracellular loop 2 (EL2) from 164-191 amino acids. It also contains two thermostabilizing mutants V74A, L415A and a F471L mutation in the vestibule to resemble human norepinephrine transporter (hNET). Additional mutations in the subsite B or subsite C were incorporated into this gene used for carrying out the uptake assays.

The uptake active dDAT construct used for co-crystallizing with norepinephrine (dDAT_mfc_) has amino acid deletions Δ1-20 and Δ162-202 along with the two thermostabilizing mutations (V74A and L415A). It also has the F471L mutation in the vestibule.

dDAT_subB_ has deletions Δ1-20 and Δ162-202; thermostabilizing mutations V74A, L415A along with two mutations in subsite-B of the substrate binding pocket D121G and S426M. This construct was used to elucidate the crystal structures of substrate-free and the duloxetine bound form.

dDAT_NET_ is identical to dDAT_subB_ with the additional mutation F471L in the vestibule. This construct was used to decipher the crystal structures in norepinephrine, milnacipran and tramadol bound complexes of dDAT.

### Expression and purification of the transporter

The recombinant expression of dDAT constructs in pEG BacMam vector was done using baculovirus mediated protein expression in mammalian cells, where HEK293S GnTI^-^ cells were transduced with high titer recombinant baculovirus using the *BacMam* method (Goehring et al., 2014). The expressed dDAT protein was extracted from membranes in 20 mM dodecyl maltoside (DDM), 4 mM cholesteryl hemisuccinate (CHS), 50 mM Tris-Cl and 150 mM NaCl. The solubilized protein was affinity purified using Talon resin in 20 mM Tris-Cl (pH 8.0), 300 mM NaCl, 100 mM imidazole containing 1 mM DDM and 0.2 mM CHS. The IMAC purified protein was treated with thrombin for removing the C-terminal GFP-8x His tag. The thrombin cleaved protein was purified by size-exclusion chromatography on a superdex-200 10/300 increase column in 4 mM decyl β D-maltoside, 0.3 mM CHS, 0.001% (w/v) 1-pamitoyl-2-oleoyl-*sn*-glycero-3-phospho ethanolamine (POPE), 20 mM Tris-Cl (pH 8.0), 300 mM NaCl and 5% glycerol. The peak fractions were collected and pooled before incubating with the substrate (6 mM) or inhibitor (0.5 – 2.0 mM) to the indicated final concentrations used for crystallization. For complexes containing norepinephrine, 4mM ascorbic acid was added to prevent its oxidation.

### Heterologous expression and purification of Fab

Fab 9D5 was cloned into pFastBac Dual vector with the heavy chain under the polyhedron promoter and the light chain under the P10 promotor with an N-terminal GP64 signal peptide on each chain. A TEV protease site followed by 8x-His tag was added to the C-terminus of heavy chain. The cloned plasmid was transformed into *E.coli* DH10Bac for the generation of recombinant bacmids. The bacmids were then transfected into Sf9 cells to generate recombinant baculovirus. High titre recombinant virus was used to infect large volume cultures and 96 hours post-infection, the cells were spun down and the supernatant containing Fab was dialysed against 25 mM Tris-Cl (pH 8.0) and 50 mM NaCl. The dialysate was then passed through Ni-NTA beads, washed with 50 mM imidazole and eluted in 25 mM Tris-Cl (pH 8.0), 50 mM NaCl containing 250 mM imidazole. The eluted Fab was further purified by size-exclusion chromatography in 25 mM Tris-Cl (pH 8.0) and 50 mM NaCl using a superdex 75 10/30 column. The purified Fab was stored at 4°C.

### Crystallization and structure determination

The SEC purified dDAT was incubated with varying concentrations of ligands for 2-4 hours at 4°C before complex formation with the recombinant antibody fragment (Fab) 9D5 in a molar ratio of 1:1.2 (dDAT:Fab) for further 30 mins on ice. The DAT-9D5 complex was concentrated to a final concentration of 3.0-4.0 mg/mL using a 100 kDa cutoff centrifugal concentrator (Amicon Ultra). The concentrated sample was clarified to remove aggregates by high-speed centrifugation at 13000 rpm for 30 mins. The clarified sample was then subjected to crystallization by hanging-drop vapor diffusion method at 4°C. Crystals of dDAT_subB_ and dDAT_NET_ proteins were obtained after 2-4 weeks, while those for dDAT_mfc_ were obtained only after a week of seeding the crystallization drops with the dDAT_NET_ crystals with a nylon fiber. All crystals were obtained in 0.1M MOPS, pH 6.5 – 7.0 and 30-32% PEG 600 as precipitant at 4°C. Data from crystals were collected at different synchrotrons sources (Table S1) and crystals for all datasets diffracted in a resolution range of 2.8-3.3 Å. The diffraction data was processed using XDS (Kabsch, 2010) and merged and scaled using AIMLESS in the CCP4 software suite (Winn et al., 2011). Five percent of the reflections were randomly assigned for R_free_ calculations as part of cross-validation. The structures were solved by molecular replacement through PHASER (McCoy et al., 2007) using the coordinates of dDAT and 9D5 from PDB ids 4XP1 or 4XNX. Refinement of the coordinates against the diffraction data was done using phenix.refine in the PHENIX crystallographic software suite (Zwart et al., 2008). Protein and inhibitor structures were built using COOT (Emsley and Cowtan, 2004) and modeling for lower resolution datasets was done with the aid of feature-enhanced map (FEM) employed in the PHENIX suite (Afonine et al., 2015).

### Whole-cell uptake inhibition assays

Whole cell-based inhibition of [^3^H] dopamine uptake was carried out in HEK293S GnTI^-^ cells transfected with pEG BacMam plasmid having dDAT CGFP gene or its mutants. Cells were washed in 1x PBS and harvested 35-40 hours post-transfection into the assay buffer (25 mM HEPES-Tris pH 7.1, 130 mM NaCl, 5.4 mM KCl, 1.2 mM CaCl_2_, 1.2 mM MgSO_4_, 1 mM ascorbic acid, 5 mM glucose and 30 μM Pargyline, a monoamine oxidase inhibitor). Whole cells were incubated with indicated concentrations of inhibitors or substrates for 20 min at room temperature. Uptake activity measured from cells incubated with 25 μM of desipramine, was used as control. Following this, 2 μM tritiated [^3^H] dopamine (DA) (1:50 or 1:100 molar ratio of [^3^H]-DA:[^1^H]-DA) was added and incubated for 20 min at room temperature. Dopamine uptake was arrested by adding 1mL of ice-cold uptake buffer containing 2 μM desipramine to each reaction followed by washing the cells twice with the same buffer to remove excess [^3^H] DA. Cells were then solubilized in 0.1 mL of 20 mM DDM and 4 mM CHS for one hour at room temperature followed by high-speed centrifugation to remove un-solubilized material. The supernatant was added to liquid scintillation fluid and the counts were estimated using a MicroBeta scintillation counter (Perkin Elmer). The background subtracted, dose-response plots were analyzed using GraphPad Prism v.5.0.1 and *K*_i_ values were determined from Cheng-Prusoff’s equation using the IC_50_ values obtained from the experiments. For *K*_M_ determination, the transfected HEK293S GnTI^-^ cells were incubated varying concentrations of DA (0.2 μM, 1 μM, 2 μM, 10 μM, 20 μM and 40 μM) in 1:200 molar ratio of [^3^H]-DA : [^1^H]-DA. The uptake was arrested at different time points (1, 2, 3 and 4 mins) with 100 μM desipramine. Initial uptake rates were used to deduce the *K*_M_ value.

### Binding and competition assays

Binding assays were performed with 10 nM of purified dDAT protein by scintillation proximity assay (SPA). For determining nisoxetine *K*_D_, [^3^H] Nisoxetine (in 1:10 molar ratio) was used in the range of 1 nM to 600 nM with 100 µM desipramine added to the control samples. Competition assays were done with 50 nM [^3^H] Nisoxetine (in 1:10 molar ratio) with the concentration range for tramadol being 10 nM to 1 mM, duloxetine from 0.1 nM to 3 µM and milnacipran from 1 nM to 100 µM. The assays were done in 1 mM DDM, 0.2 mM CHS, 20 mM Tris-Cl (pH 8), 300 mM NaCl and 5% glycerol. The background values were subtracted to plot the final curves in GraphPad prism v5.0.1 and the *K*_D_ values were calculated. The IC_50_ values obtained from binding competition assays were used to deduce the *K*_*i*_ values by Cheng-Prusoff’s equation.

### Data resources

The coordinates for the structures have been deposited in the Protein Data Bank with the following accession codes 6XX1, 6XX2, 6XX3, 6XX4, 6XX5, 6XX6.

