## Supplementary Table and Figures for "Structural basis of norepinephrine recognition and transport inhibition in neurotransmitter transporters"

### Table S1. Data collection and Refinement statistics

| Parameter | Substrate-free dDAT$_{subB}$ | NE-dDAT$_{mfc}$ | NE-dDAT$_{NET}$ | S-duloxetine dDAT$_{subB}$ | Milnacipran dDAT$_{NET}$ | Tramadol dDAT$_{NET}$ |
|---|---|---|---|---|---|---|
| **Source** | ESRF-ID29 | ElettraXRD2 | APS-24IDC | ElettraXRD2 | APS-24IDC | SLS-PX1 |
| **Wavelength** | 0.968 | 0.979 | 0.979 | 0.979 | 0.979 | 1.000 |
| **Space group** | $P2_12_12_1$ | $P2_12_12_1$ | $P2_12_12_1$ | $P2_12_12_1$ | $P2_12_12_1$ | $P2_12_12_1$ |
| **Cell dimensions** ***a, b, c*** **(Å);** **α = β = γ = 90°** | 97.7, 133.3, 162.0 | 96.8, 141.2, 167.6 | 97.1, 142.5, 168.0 | 97.3, 140.6, 167.8 | 96.5, 140.9, 167.3 | 96.8, 140.8, 167.6 |
| **Resolution (Å), (*)** | 50.0-3.3 (3.5-3.3) | 50.0-2.8 (2.97-2.8) | 50.0-2.88 (3.05-2.88) | 50.0-3.0 (3.18-3.0) | 50.0-3.11 (3.29-3.11) | 50.0-3.25 (3.45-3.25) |
| **Total Observations** | 180963 (27837) | 352419 (55239) | 358042 (55889) | 338525 (54597) | 228053 (37703) | 193872 (32191) |
| **Unique reflections** | 32633 (5148) | 109219 (17635) | 53971 (8591) | 88712 (14205) | 42395 (6747) | 64838 (10759) |
| **Multiplicity** | 5.5 (5.4) | 3.22 (3.13) | 6.6 (6.5) | 3.81(3.84) | 5.4(5.6) | 3.0 (3.0) |
| **Completion (%)** | 99.7(99.0) | 99.5 (98.9) | 99.9 (99.9) | 99.6 (99.0) | 99.6(99.3) | 93 (95.8) |
| **I/σI** | 10.6(0.94) | 11.4 (1.1) | 13.4(1.0) | 11.3 (1.2) | 7.64(1.0) | 8.39 (1.0) |
| **$CC_{1/2}$ (%)** | 99.9(60.9) | 99.8 (53.6) | 99.8 (43.3) | 99.8 (49.3) | 99.7 (47.5) | 99.9 (84.1) |
| **$R_{meas}$ (%)** | 11.6 (148.9) | 10.3 (119.0) | 10.2 (206.6) | 10.7 (116.2) | 17.4 (135.4) | 15.1 (132.3) |
| ***Refinement Statistics*** | | | | | | |
| **Resolution** | 50.0-3.3 | 50.0-2.8 | 50.0-2.88 | 50.0-3.0 | 50.0-3.1 | 50.0-3.25 |
| **Reflections#** | 32557 (3144) | 57114 (5587) | 53940 (5335) | 46691 (4542) | 42341 (4157) | 35194 (3543) |
| **$R_{work}$ (%)** | 24.3 (37.0) | 21.2 (34.3) | 21.2 (34.0) | 21.5 (30.9) | 22.3 (32.3) | 24.2 (32.2) |
| **$R_{free}$ (%)** | 27.6 (40.5) | 24.8 (39.0) | 24.7 (36.4) | 24.4 (32.5) | 26.5 (36.8) | 27.8 (37.3) |
| **No. of non-H atoms** | 7508 | 7780 | 7654 | 7634 | 7670 | 7615 |
| Protein | 7430 | 7538 | 7507 | 7490 | 7504 | 7455 |
| Substrate/Inhibitor | - | 12 | 12 | 21 | 18 | 19 |
| Ions/Lipid/Detergent | 70 | 131 | 99 | 99 | 128 | 122 |
| water | 8 | 99 | 36 | 24 | 20 | 19 |
| **Average B-factor** | 113.6 | 64.2 | 89.92 | 78.3 | 84.6 | 89.9 |
| Protein | 113.6 | 64.05 | 89.73 | 78.2 | 84.4 | 89.5 |
| Substrate/Inhibitor | - | 87.8 | 102 | 71.5 | 79.7 | 88.0 |
| Detergent/Lipid | 116.0 | 77.4 | 101 | 89 | 97 | 101.4 |
| water | 93.1 | 60.1 | 92.7 | 69.9 | 75.8 | 80.1 |
| **RMS bond lengths (Å)** | 0.004 | 0.006 | 0.005 | 0.006 | 0.01 | 0.005 |
| **RMS bond angles (°)** | 1.08 | 1.12 | 1.00 | 1.1 | 1.40 | 1.17 |
| **Ramachandran statistics** | | | | | | |
| Favoured | 92.2 | 94.60 | 94.70 | 95.8 | 94.8 | 95.1 |
| Allowed | 7.8 | 5.4 | 5.3 | 4.2 | 5.2 | 4.9 |
| Disallowed | 0 | 0 | 0 | 0 | 0 | 0 |

*** Values in parentheses represent highest-resolution shell.**

**# Reflections employed in structure refinement.**

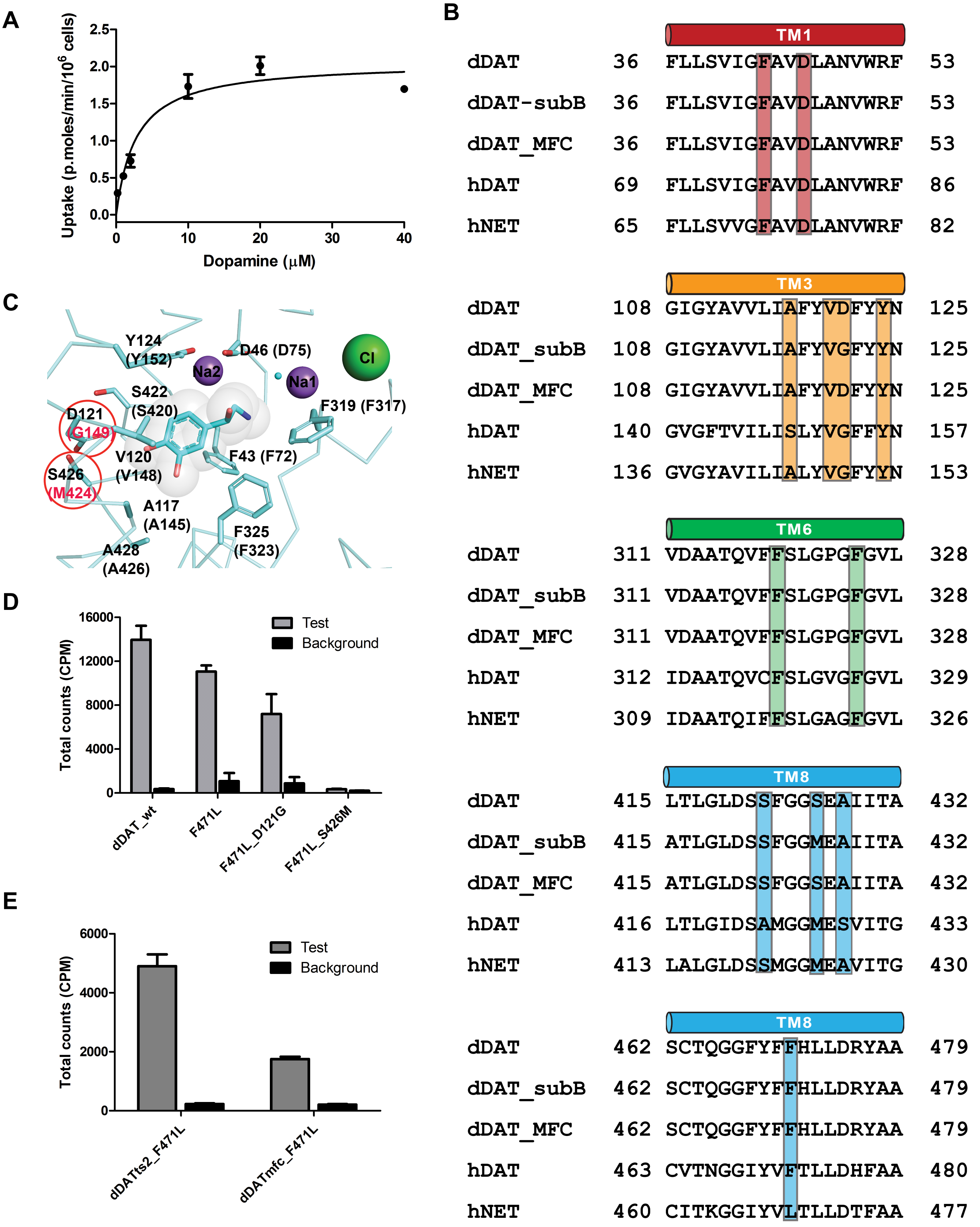


**Figure S1. (A)** [$^3$H] Dopamine uptake kinetics of dDAT functional construct with F471L mutation. Graph represents one experiment out of three trials, each performed in duplicate. $K_M$ value measured was 4.5 ± 2.4 μM. **(B)** Multiple sequence alignment of the helices surrounding the primary substrate/inhibitor binding site with residues that sculpt the binding pocket highlighted as vertical bars. **(C)** Binding site residues and subsiteB mutants performed in the study represented in the structure with bound L-norepinephrine. **(D)** [$^3$H] Dopamine uptake analysis using subsiteB mutants D121G and S426M in the context of F471L mutant. S426M causes a loss of transport activity. **(E)** Uptake activity of $dDAT_{mfc}$ carrying F471L done in comparison with $dDAT_{WT}$ with F471L and two thermostable mutants. Experiments represent one trial performed in duplicates.

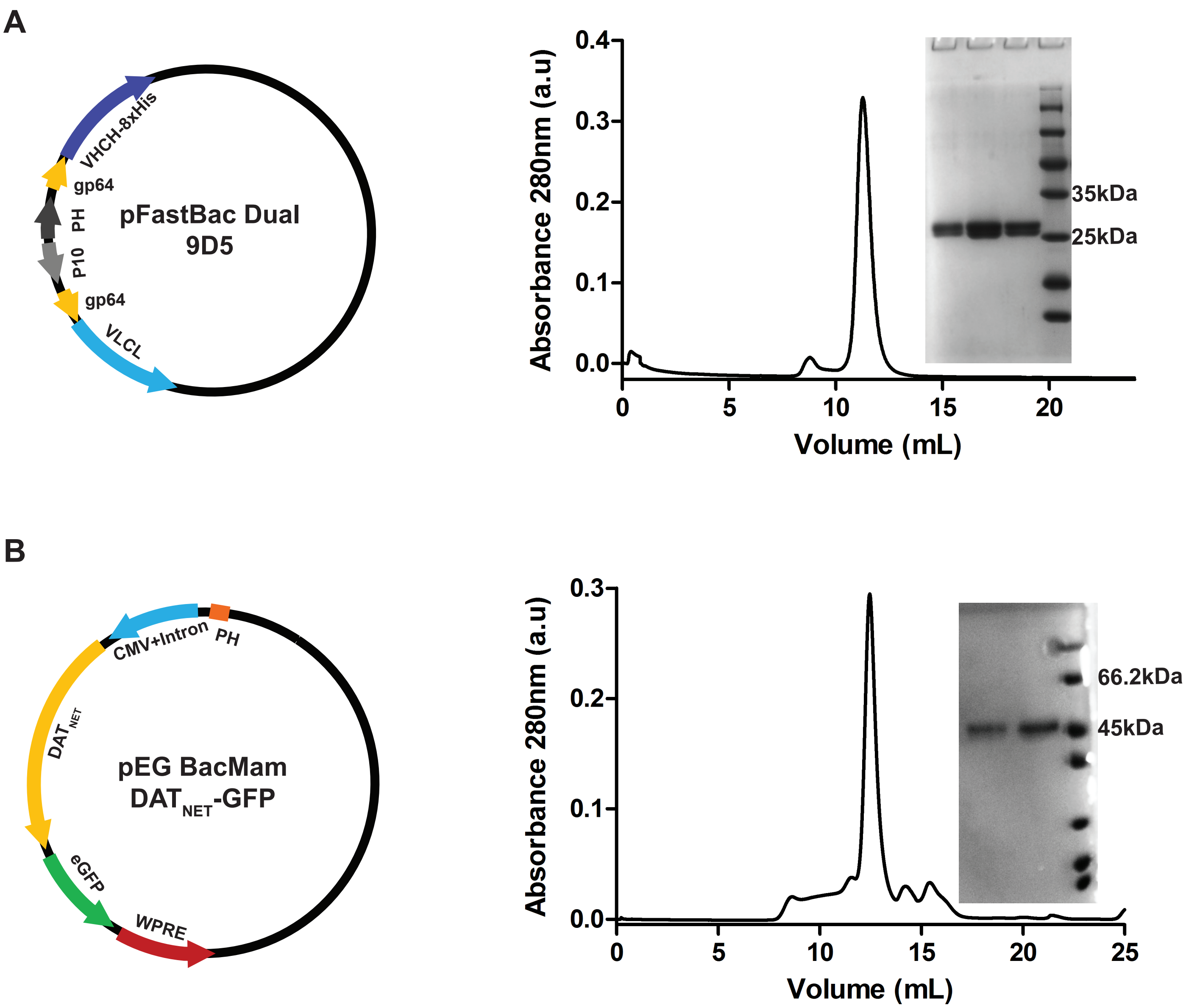


**Figure S2. (A)** Vector design of pFastBac dual containing the heavy and light chains of the 9D5 antibody fragment cloned downstream of the polyhedrin (PH) and p10 promoters respectively. Both genes were cloned with a gp64 signal peptide at the N-terminus to facilitate export from insect cells. Panel on the right shows size exclusion chromatogram of the purified Fab and inset displays heavy and light chain bands for the purified 9D5 with coomassie stained SDS-PAGE (12%). **(B)** Vector design of the dDAT constructs in pEG Bacmam with a C-terminal GFP that was cleaved by thrombin cleavage post-expression. Panel on right shows chromatogram of purified dDAT and the corresponding protein band stained with coomassie on an SDS-PAGE (12%).

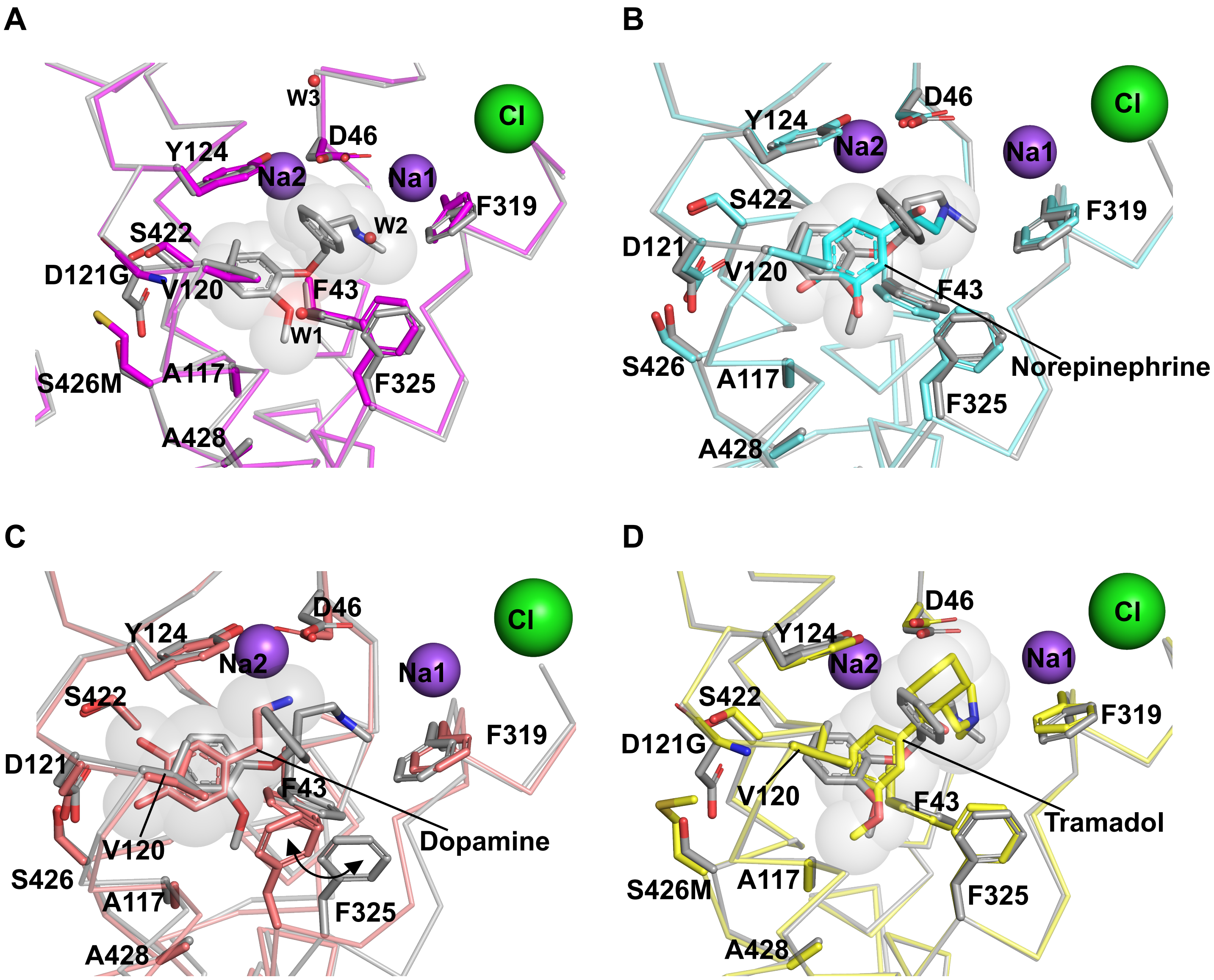


**Figure S3. (A)** Overlay of nisoxetine bound dDAT (gray) (4XNU) with substrate-free conformation of $dDAT_{subB}$ to highlight differences in the binding pocket. Similar structural comparisons with **(B)** L-nor-epinephrine bound $dDAT_{mfc}$, **(C)** dopamine bound $dDAT_{mfc}$, **(D)** tramadol bound $dDAT_{NET}$ in the primary binding site.

### Substrate/Inhibitor electron densities

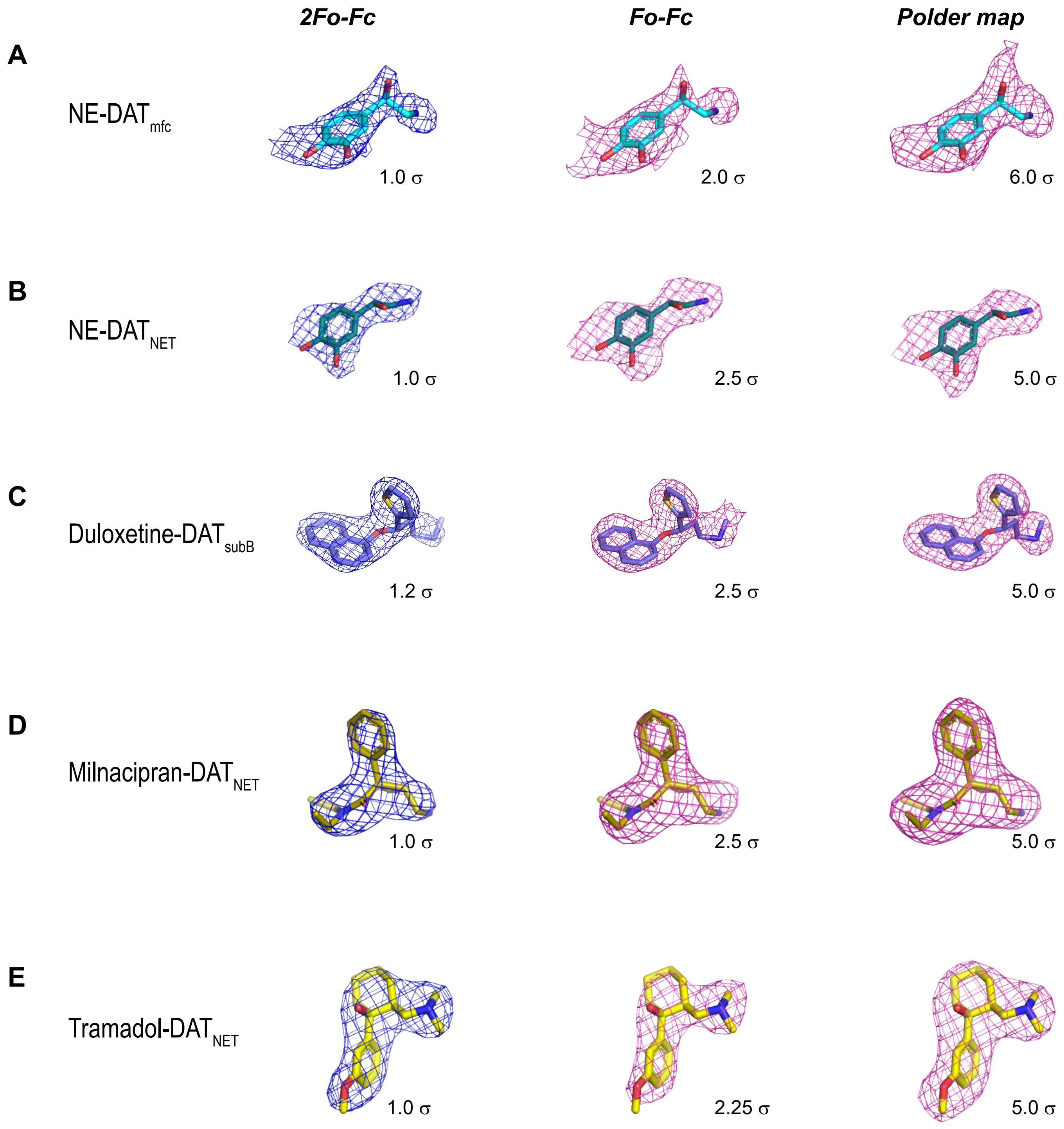


**Figure S4.** Electron densities of substrate and inhbitors bound in the binding pocket of dDAT structures in this study. Column 1 represents 2*Fo-Fc* maps (blue), column 2 represents *Fo-Fc* omit maps (magenta) and column 3 represents Polder maps (magenta) for **(A)** NE bound $dDAT_{mfc}$ **(B)** NE bound $dDAT_{NET}$ **(C)** duloxetine bound $dDAT_{subB}$ **(D)** milnacipran bound $dDAT_{NET}$ **(E)** tramadol bound $dDAT_{NET}$.

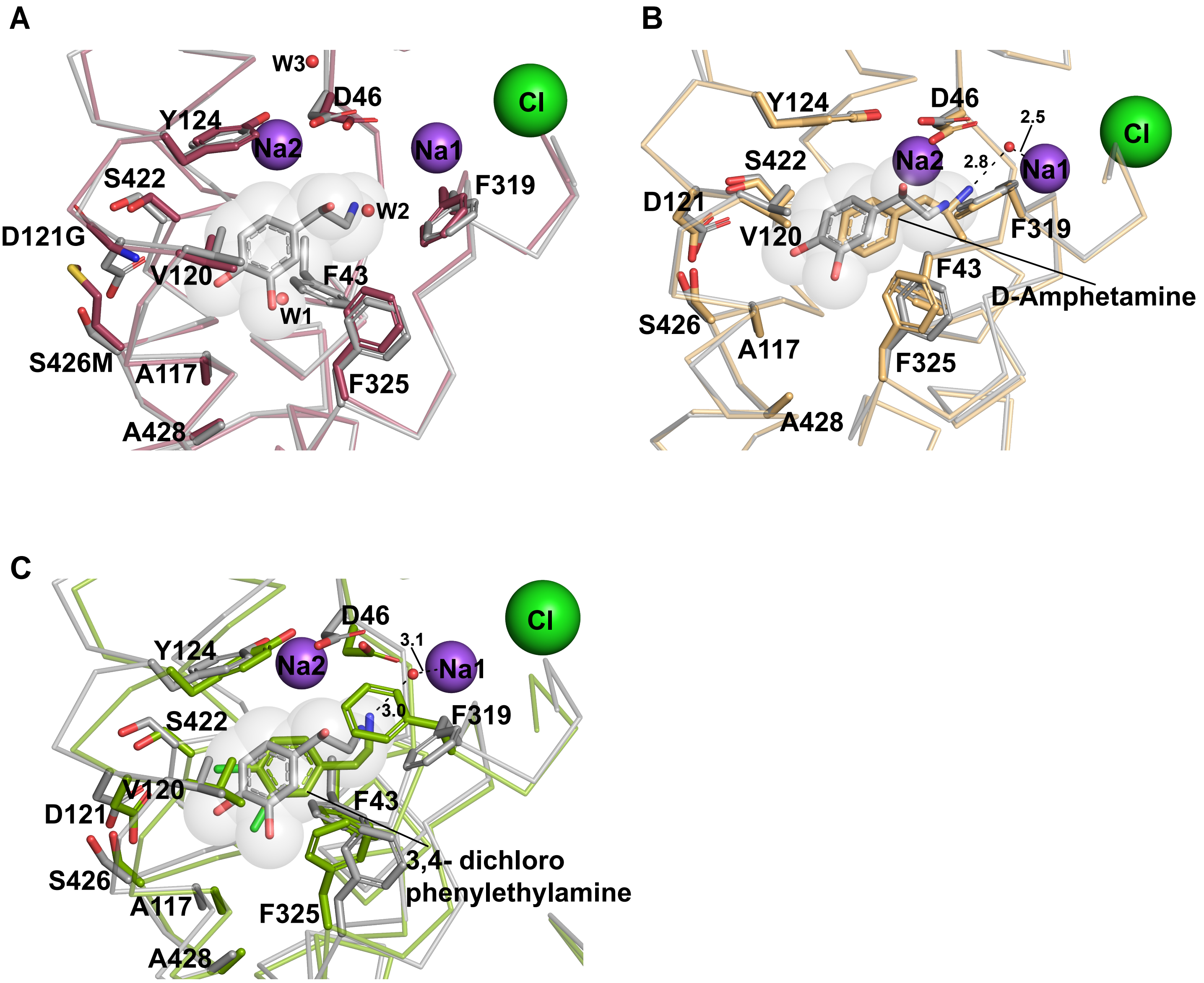


**Figure S5**. Structural comparisons between NE bound $dDAT_{mfc}$ (gray) with **(A)** substrate free state of dDAT, **(B)** D-amphetamine bound $dDAT_{mfc}$ (4XP9 ), **(C)** 3,4 dichlorophenethylamine bound $dDAT_{mfc}$ (4XPA).

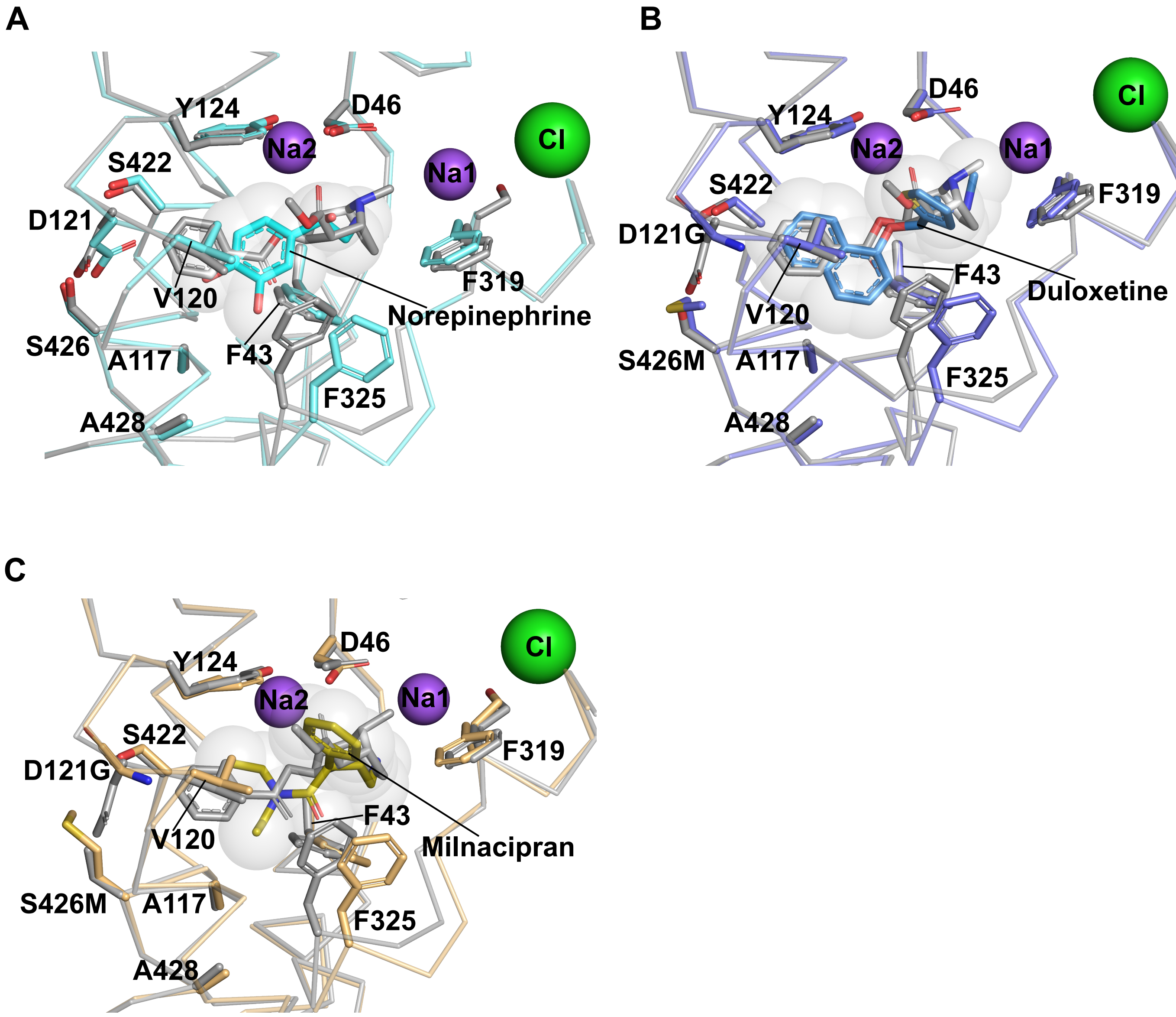


**Figure S6**. Structural comparisons between cocaine bound dDAT$_{mfc}$ (gray) (4XP4) with **(A)** NE bound dDAT$_{mfc}$, **(B)** duloxetine bound dDAT$_{subB}$, **(C)** milnacipran dDAT$_{NET}$.

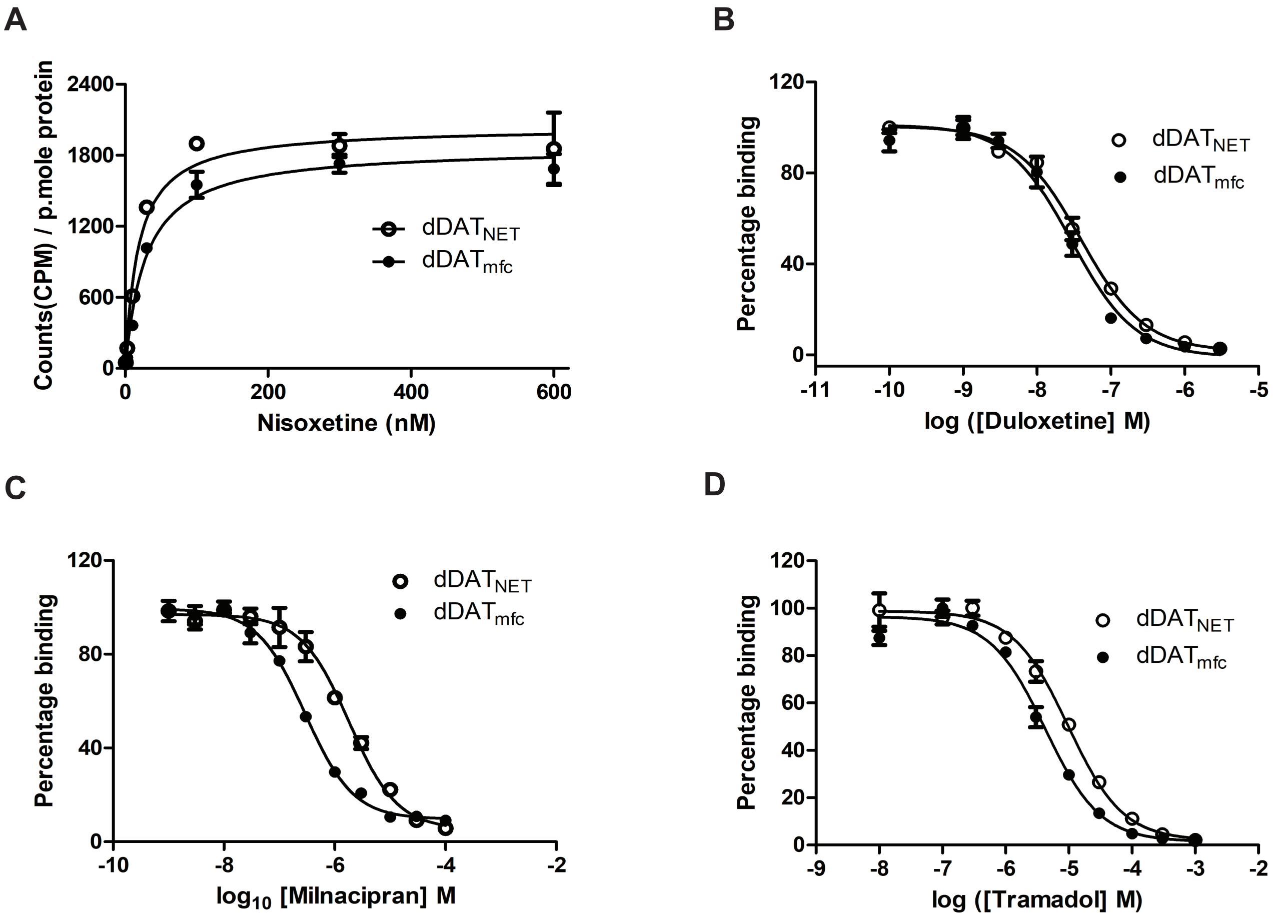


| NET inhibitor | Binding $K_i$ with dDAT$_{NET}$ | Binding $K_i$ with dDAT$_{mfc}$ |
|---|---|---|
| Duloxetine | 10.2 nM | 10.8 nM |
| Milnacipran | 480 nM | 65 nM |
| Tramadol | 2.6 μM | 1.6 μM |

**Figure S7. (A)** Binding curves of $[^3H]$ nisoxetine with dDAT$_{mfc}$ and dDAT$_{NET}$ containing mutations in primary binding site D121G and S426M. Graph shown is one of four independent experiments each time performed in duplicate. Error bars at each point represent the range of counts observed at a given concentration of $[^3H]$ nisoxetine. $K_d$ values of 28.7± 4.3 nM and 18.4 ± 3.9 nM were measured for dDAT$_{mfc}$ and dDAT$_{NET}$ respectively. Competitive inhibition of $[^3H]$ nisoxetine binding with increasing concentration of **(B)** duloxetine, **(C)** milnacipran, **(D)** tramadol performed for dDAT$_{mfc}$ (solid circles) and dDAT$_{NET}$ (open circles). All curves are one of two trials done each time in triplicate. $K_i$ values reported in table below were calculated from estimated $IC_{50}$ values of each inhibition curve and $K_d$ values estimated from nisoxetine binding using the Cheng-Prusoff equation.
